# ALKBH1 drives codon-biased, pro-oncogenic translation and tumor microenvironment remodeling in glioma via tRNA wobble oxidation

**DOI:** 10.1101/2025.11.28.691056

**Authors:** Atsushi Nakayashiki, Arata Nagai, Yuko Iwasaki, Abdulrahman Mousa, Moe Kumai, Munehiro Kimura, Shadi Al-Mesitef, Hiroshi Kanno, Shota Yamashita, Daisuke Ando, Yoshiteru Shimoda, Masayuki Kanamori, Keisuke Goda, Hidenori Endo, Sherif Rashad, Kuniyasu Niizuma

## Abstract

Gliomas rely on translational plasticity to sustain heterogeneity, stemness, and immune evasion. Here we identify the tRNA dioxygenase ALKBH1 as a central regulator of codon-biased translation and tumor microenvironment remodeling. LC-MS/MS profiling of patient tumors revealed enrichment of ALKBH1-mediated wobble cytidine oxidation modifications in high-grade gliomas. Genetic perturbation demonstrated that ALKBH1 overexpression slows proliferation in vitro but worsens survival in vivo by promoting glioma stem-like and neuronal states and suppressing anti-tumor immunity. Ribosome profiling showed that ALKBH1 establishes an A/T-ending codon-biased translational program, enhancing decoding of rare leucine codons (TTA/TTG) and driving synthesis of pro-stemness transcripts. Single-cell RNA-seq analyses further revealed that ALKBH1 promotes glioma heterogeneity and induces neuronal cell clusters and rewires intercellular communication ECM-related signaling while dampening immune pathways and reducing immune infiltration. These findings establish a mechanistic link between tRNA oxidation, codon bias, and glioma aggressiveness, positioning ALKBH1 as a potential therapeutic target for glioma.

## INTRODUCTION

Glioblastoma (GB) is the most common and deadliest adult primary brain tumor^1,2^. Five-year survival remains poor, and survivors often experience substantial neurologic morbidity and impaired quality of life because of the tumor’s involvement of eloquent brain networks^1^. GB accounts for approximately half of malignant primary brain tumors, whereas diffuse infiltrating low-grade gliomas (LGG) constitute about 30%^1^. Adult diffuse gliomas comprise three molecularly and prognostically distinct entities: IDH-mutant astrocytoma, IDH-mutant oligodendroglioma, and IDH-wild-type glioblastoma. Standard-of-care therapy (i.e., maximal safe resection followed by radiotherapy and temozolomide) offers limited durable control^1,2^. Despite the success of immunotherapies and targeted agents in other solid tumors, outcomes in GB remain dismal^3^. Several features contribute to this therapeutic recalcitrance: marked intratumoral heterogeneity and plasticity driven in part by glioma stem-like cells^4^; the ability of GB to evade, suppress, and even co-opt immune responses^5,6^; and extensive crosstalk with neurons and glia that rewires circuit activity to support tumor growth^7,8^. While genomic and transcriptomic landscapes have been deeply profiled, the regulatory layers that operate beyond DNA sequence, particularly at the epitranscriptomic and translational levels, are less well understood but are likely central to phenotypic switching, treatment resistance, and microenvironmental adaptation.

Transfer RNA (tRNA) modifications are emerging as critical regulators of translational control in cancer^9^. On average, each tRNA harbors roughly a dozen modifications^10,11^. Chemical changes at the anticodon wobble position (nucleotide 34) can broaden or restrict pairing with synonymous codons and thereby redistribute translational efficiency across codon families^9–14^. Numerous tRNA-modifying enzymes are dysregulated in malignancy, including glioma^9,11,15^. For example, elevated 7-methylguanosine (m^7^G) modification, driven in part by copy-number gains of the methyltransferase METTL1, supports an oncogenic translational program in several cancers^9,16–18^. In GB, N6-threonylcarbamoyladenosine (t^6^A) promotes tumor growth via codon-biased translation^15^, and 2-methylthio-N6-isopentenyladenosine (ms^2^i^6^A) helps maintain glioma stemness and self-renewal^19^. These studies, together with work in other malignancies, indicate that tRNA chemistry can be tuned to favor specific codon pools and cell states. Nonetheless, key gaps remain. First, how wobble-site oxidation and editing are deployed to shape codon usage in glioma is not resolved. Second, expression of the “writers,” “erasers,” and “readers” of tRNA marks often correlates poorly with the stoichiometry of the modifications themselves, which also depends on tRNA transcription, turnover, and interdependent modification cascades^11,12,20^. Consequently, enzyme-centric analyses are insufficient; direct, modification-centric profiling is required to define the epitranscriptomic architecture of glioma.

In this study, we apply a modification-centric framework to human gliomas and mechanistically delineate how a tRNA dioxygenase couples codon bias to malignant phenotypes. Using LC-MS/MS, we quantify tRNA modifications in surgically resected gliomas spanning grades and molecular classes and find that wobble-site oxidation marks, namely 5-hydroxymethylcytidine (hm5C) and 5-formylcytidine (f5C), and their 2’-O-methylated derivatives are significantly enriched in high-grade tumors (Grade IV astrocytoma and glioblastoma) irrespective of IDH status. We then focus on ALKBH1, a tRNA dioxygenase/demethylase implicated in wobble cytidine oxidation^21–25^. We show how ALKBH1 drives a codon biased pro-oncogenic translational program that promotes the translation of rare leucine codons. Through this translational program, ALKBH1 drives a pro-stemness pro-neuronal phenotype in glioma that allows it to evade immune detection, sustain heterogenous cell populations, and remodel its microenvironment to sustain a more aggressive phenotype.

## RESULTS

### Analysis of the tRNA modification landscape in glioma

We began our investigation into the potential role of tRNA modifications in glioma by asking whether specific modifications correlate with tumor grade or genetic background. To address this, we collected surgical glioma specimens from patients at Tohoku University Hospital, including glioblastoma (Grade 4, GB; N = 11), Grade 2/3 astrocytoma (N = 5; low grade astrocytoma), Grade 4 astrocytoma (N = 4; high grade astrocytoma), and Grade 2/3 oligodendroglioma (N = 9). From each tumor, we isolated tRNA-enriched small RNA fraction (>90% tRNA), digested it to mononucleosides, and performed targeted LC–MS/MS analysis of RNA/tRNA modifications (Figure 1a). This revealed a global enrichment of tRNA modifications in high-grade gliomas (HGG; GB and Grade IV astrocytoma) compared with low-grade gliomas (LGG), with only a few exceptions (Figure 1b-c; Supplementary Table 1). Importantly, this enrichment followed tumor grade rather than IDH genotype: IDH-mutant high-grade astrocytomas clustered with IDH-wild-type GB (Figure 1b-c), suggesting that tRNA modification patterns reflect malignant potential more than genetic subtype. Of the 38 detected modifications, 26 were statistically significant by ANOVA (*p* < 0.05), prompting further analysis of which modifications drove the separation among tumor groups.

**Figure 1.**
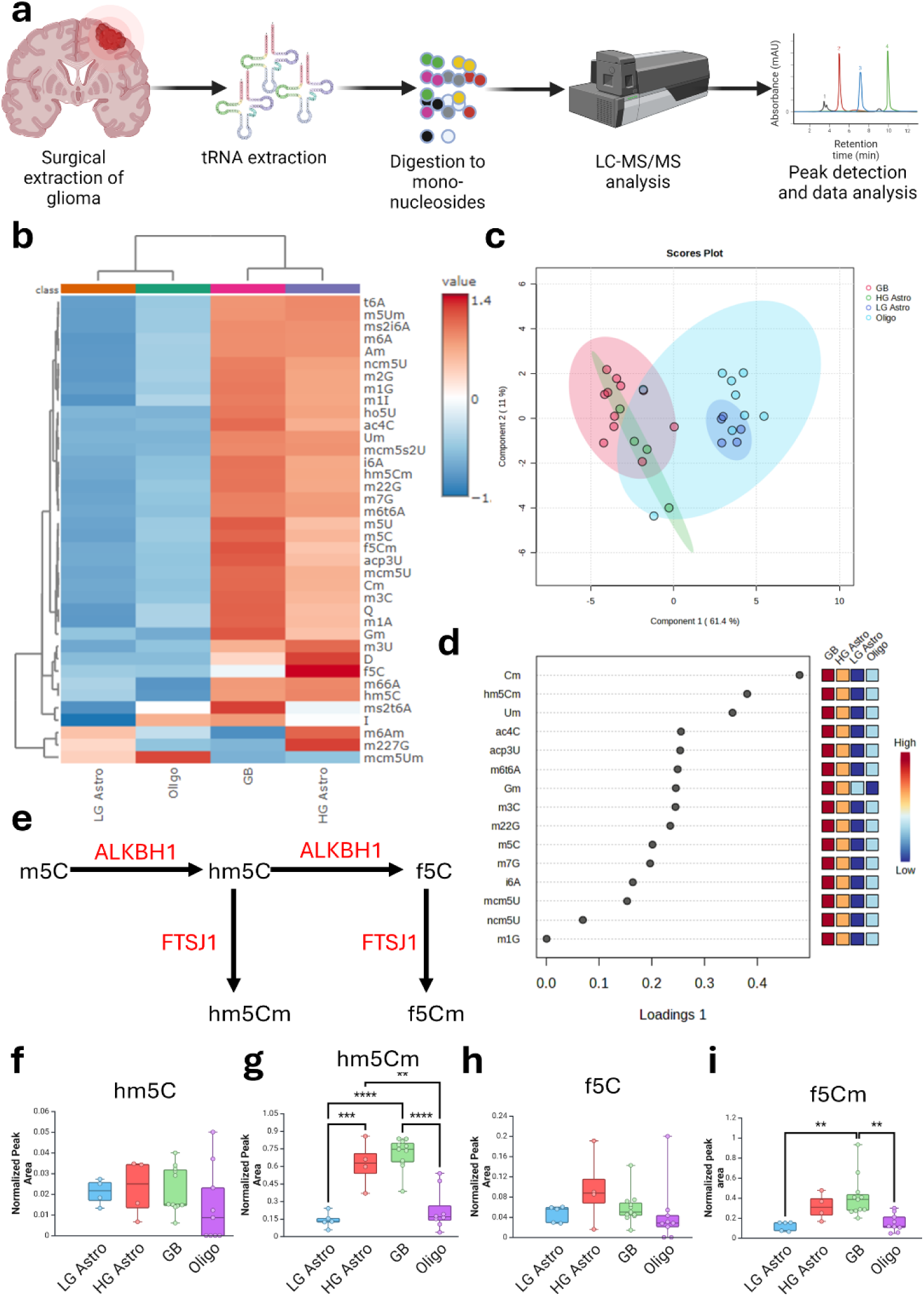
Analysis of tRNA modifications in glioma surgical samples. **a**, Methodology overview of the analytical approach. **b,** Heatmap showing the normalized expression of the detected tRNA modifications in glioma groups. Heatmap coloring shows row z-score. **c,** Sparse least square regression discriminant analysis (sPLS-DA) clustering of the glioma samples and groups based on tRNA modifications levels. **d,** Variables of importance in projection (VIP) analysis of the sPLS-DA clustering showing which modifications are important for the observed clustering patterns. **e,** The ALKBH1/FTSJ1 modifications system. **f-i,** Box plots of the selected ALKBH1/FTSJ1 modifications expression in different glioma groups. ANOVA with Turkey’s post-hoc was conducted to evaluate significance. **: *p* < 0.005, ***: *p <* 0.0005, ****: *p* < 0.0001.

To identify the most influential contributors, we performed partial least square determinant analysis (PLS-DA) followed by VIP (variable importance in projection) analysis (Figure 1c-d). Several 2’O-ribose methylation modifications, including Cm, hm^5^Cm, and Um, were major discriminators (full names in Supplementary Table 2). In addition, modifications previously associated with oncogenesis were elevated in HGGs^26,27^: ac4C (found in both mRNA and tRNA^27^), m^7^G (one of the most extensively studied tRNA cancer-associated modifications^9^), and t^6^A along with its derivatives m^6^t^6^A and ms2t^6^A, consistent with recent findings implicating t^6^A as a glioma oncogenic driver^15^. Given the large number of upregulated modifications, we focused specifically on those that directly regulate codon-anticodon interactions, especially wobble-position (position 34) modifications^10,11^, as they can strongly influence codon-biased translation, decoding efficiency, and downstream oncogenic phenotypes^9,11,28,29^. Among anticodon-associated modifications, hm^5^Cm was one of the highest-scoring marks in VIP analysis. This modification arises through a multistep oxidation pathway: ALKBH1 converts m^5^C to hm^5^C then to f^5^C using α-ketoglutarate and O₂, followed by 2’O-methylation by FTSJ1^11,21^ (Figure 1e). Consistent with this pathway, both hm^5^Cm and f^5^Cm were significantly elevated in HGG versus LGG (Figure 1f-i), supporting the idea that ALKBH1-linked wobble oxidation is selectively enhanced in aggressive gliomas.

To understand how enzyme expression relates to these modification patterns, we examined ALKBH1 and FTSJ1 mRNA levels in gliomas using TCGA datasets accessed via the Human Protein Atlas^30^ and UCSC Xena^31^. Surprisingly, higher expression of either enzyme correlated with better prognosis or showed no survival impact (Supplementary Figure 1), a finding paradoxical to the elevated modification levels observed in high-grade tumors. This discrepancy highlights a key challenge in epitranscriptomic analysis: enzyme expression is a poor proxy for modification abundance, which is influenced by tRNA transcription, turnover, substrate availability, and modification stoichiometry. Because FTSJ1 is not specific to anticodon modifications, and could act as a 2’O-ribose methyltransferase on other sites in the tRNA unrelated to decoding, such as the D- and T-loops, we focused subsequent mechanistic studies on ALKBH1, the upstream dioxygenase responsible for initiating wobble-position m^5^C oxidation and the most biologically specific entry point into this modification pathway^21,32^.

### ALKBH1 is a tRNA dioxygenase in glioma cells

The wobble modifications hm^5^C and f^5^C expand the decoding capacity of tRNA-Leu-CAA from the canonical Leu-UUG codon to both Leu-UUG and Leu-UUA codons^21^, thereby broadening leucine codon recognition in a manner that could influence translational programs relevant to glioma oncogenesis. To investigate how ALKBH1-mediated wobble oxidation affects glioma biology, we generated ALKBH1 knockout (KO) and constitutive overexpression (OE) models in four glioma cell lines: the rodent line 9L/GS9L, used for animal studies, and three human cell lines (U87, U251, A172). KO lines were created using CRISPR/Cas9 with scramble sgRNA controls, whereas OE lines were produced through stable lentiviral transduction alongside empty-vector controls (Figure 2a). Western blotting confirmed successful ALKBH1 depletion or overexpression in each line (Figure 2b; Supplementary Figure 2). We next quantified ALKBH1-linked tRNA modifications, such as hm^5^C, hm^5^Cm, f^5^C, f^5^Cm, and m^1^A^21,24^, to determine whether genetic manipulation altered their abundance. LC-MS/MS analysis revealed a consistent and substantial reduction of hm^5^C, hm^5^Cm, and f^5^Cm in all KO cell lines except U251 (Figure 2c-d; Supplementary Figures 2–6). We did not, on the other hand, observe significant changes in f^5^C levels after ALKBH1 KO in any of the tested cell lines. Conversely, ALKBH1 OE increased these modifications to varying degrees depending on the cell line: U87, 9L, and U251 all showed upregulation of hm^5^C and f^5^Cm, whereas f^5^C itself increased significantly only in U251.

**Figure 2.**
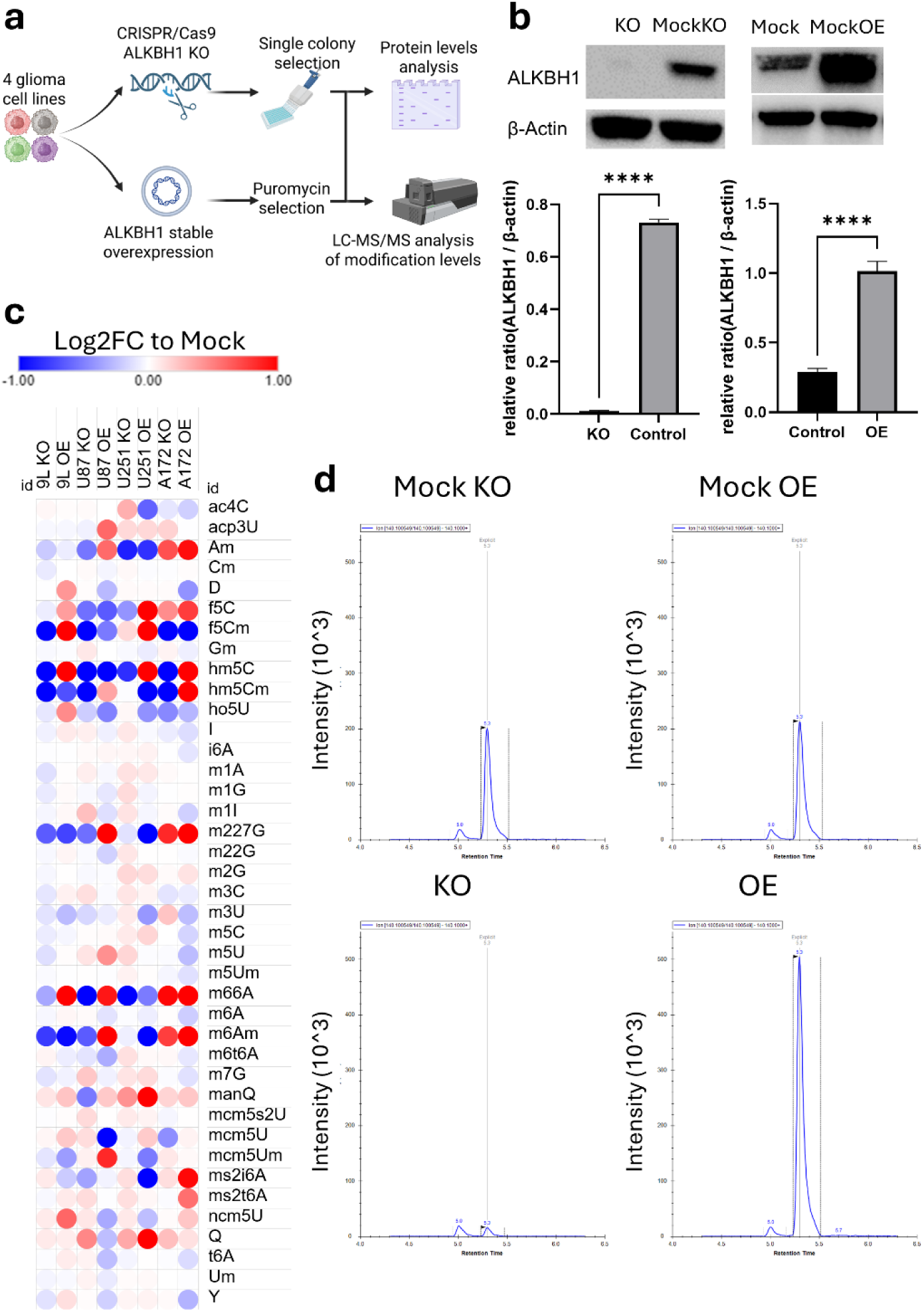
Validating ALKBH1’s role in modifying tRNA. **a**, Schematic overview of the methodology for generating and validating ALKBH1 transgenic cells. **b,** Western blot analysis of ALKBH1 expression after knockout (KO) or overexpression (OE) in GS-9L cells. Graphs show the quantification results of western blotting from 3 biological replicates. Student’s T-test was used for analyzing statistical significance. Supplementary Figure 2 shows the western blots of the other used cell lines **c,** Heatmap showing the log2 fold change values of all detected tRNA modifications compared to the appropriate Mock controls across all tested cell lines. **d,** An example chromatogram showing f^5^Cm levels in GS-9L cells. Y-axis are fixed across the 4 chromatograms to show the evident upregulation of f^5^Cm in OE cells.

These variations suggest that ALKBH1 activity exhibits context dependence, potentially reflecting differences in substrate availability, such as intracellular 2-oxoglutarate (α-ketoglutarate)^21^, a required cofactor for its dioxygenase reaction, or cell-specific tRNA transcription and turnover^24^. Our data also hints at the potential presence of another unidentified dioxygenase given that f^5^C levels remained unchanged after ALKBH1 KO. This ALKBH1-independent tRNA oxidation was previously observed in cytosolic tRNAs Val-CAC, Gly-CCC and Gln-CTG^22^. It is important to note that in HEK293T cells, we observed a stronger downregulation of f^5^C after ALKBH1 KO^12^, which could indicate that the observed phenomena are cell context specific. Such observations warrant additional mechanistic studies beyond the scope of the present work. Because ALKBH1 has also been reported to function as an m^1^A demethylase, we examined m^1^A abundance across KO and OE models; however, ALKBH1 loss did not alter m^1^A levels in any cell line tested, in agreement with prior observations that its demethylase activity may be restricted to specific contexts or mitochondrial tRNAs or during specific contexts such as oxidative stress^22,24,25^. Taken together, our data demonstrate that in glioma cells ALKBH1 primarily acts as a tRNA wobble dioxygenase, catalyzing the oxidation of m^5^C at position 34 to hm^5^C and subsequently to f^5^C^21^, thereby establishing the biochemical foundation for downstream impacts on codon decoding and translational regulation.

### ALKBH1 slows proliferation *in vitro* but worsens survival *in vivo*

We next examined how ALKBH1 influences glioma cell phenotypes in vitro and in vivo. Across all four glioma cell lines, ALKBH1 OE consistently slowed cell proliferation, whereas its KO accelerated proliferation in every line except A172 (Figure 3a-b; Supplementary Figure 7a-f). To evaluate whether these effects translated into differences in tumor behavior in vivo, we implanted transgenic 9L cells (i.e., KO, OE, and matched controls) orthotopically into Fischer 344 rats. Paradoxically, in contrast to the proliferation trends observed in vitro, ALKBH1 KO markedly improved animal survival, while OE significantly worsened it (Figure 3c-d). Tumor-size measurements revealed that KO tumors were smaller, whereas OE tumors were comparable in size to their mock controls (Figure 3e-f), indicating that differences in outcome were not simply attributable to bulk tumor growth.

**Figure 3.**
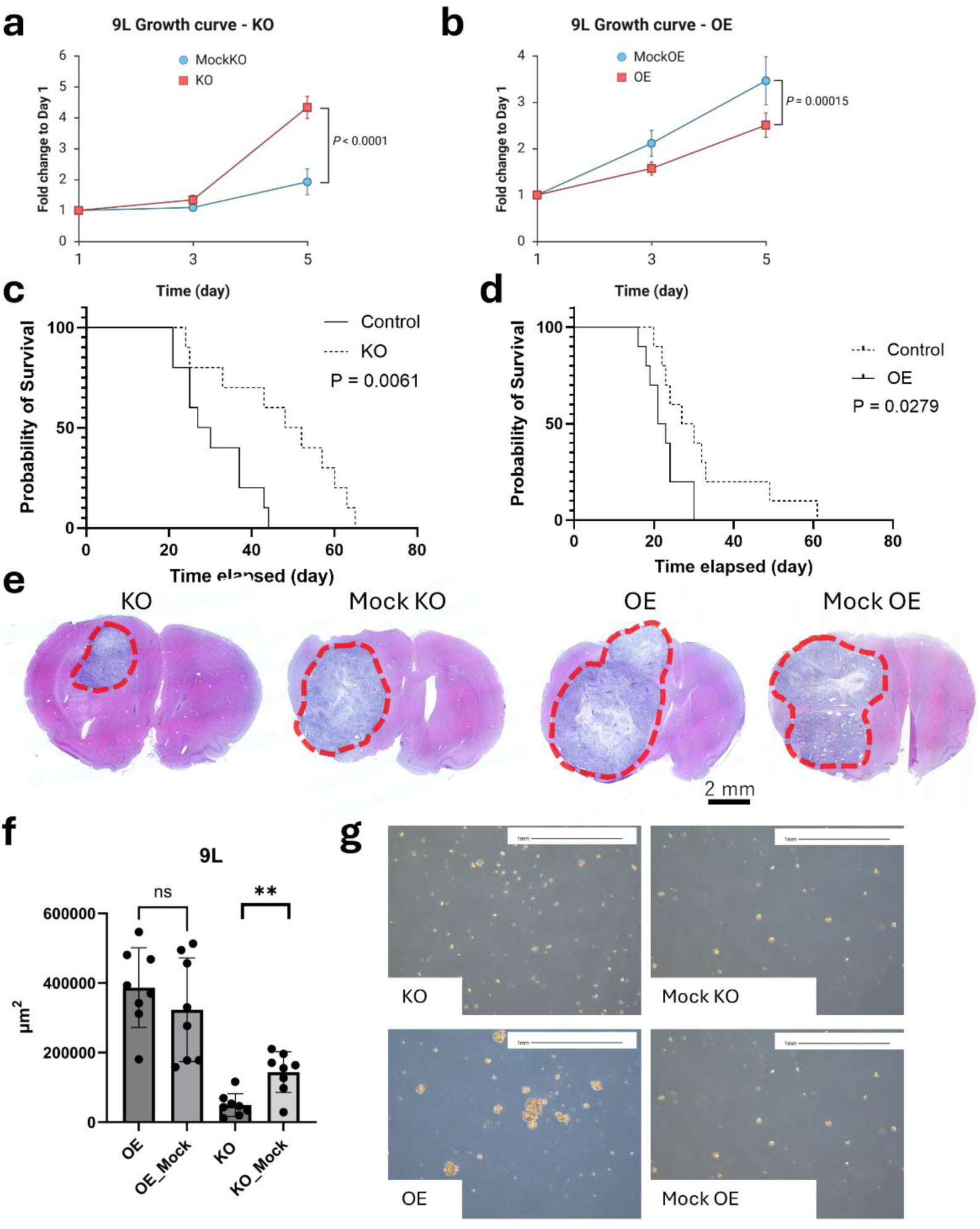
ALKBH1 reduces glioma proliferation *in vitro* but promotes oncogenesis *in vivo*. **a, b**, Analysis of GS-9L transgenic cells proliferation. Linear regression was used for statistical analysis. **c,** Kaplan-Meier analysis of Fisher rat survival after Alkbh1 KO or MockKO inoculation (N = 10 animals per group). **d,** Kaplan-Meier analysis of Fisher rat survival after Alkbh1 OE or MockOE inoculation (N = 10 animals per group, Statistical analysis was performed using Logrank (Mantel-Cox) test. **e,** Representative H&E sections showing the tumor sizes in after 21 days of inoculating transgenic GS-9L cells in Fisher 344 rats. **f,** Bar plot showing the size (in µm^3^) of tumors after 21 days of inoculation. Unpaired T-test (comparing KO to MockKO and OE to MockOE) was used for statistical analysis. **g,** GS-9L cells cultured in neurobasal medium showing spontaneous spheroid formation in Alkbh1 overexpressing cells.

In exploring the cellular basis of these effects, we found that ALKBH1 OE cells spontaneously formed spheroids when cultured in neurobasal medium, a hallmark of glioma stem-like cells, whereas neither mock nor KO cells displayed this behavior (Figure 3g). This observation suggested that ALKBH1 may promote dedifferentiation or reinforce stemness programs. Together, these findings demonstrate that ALKBH1 overexpression promotes stem-like phenotypes and worsens in vivo outcomes despite slowing proliferation in vitro, highlighting the importance of cell-state transitions rather than simple growth rate in determining glioma aggressiveness.

### ALKBH1 maintains glioma mitochondrial function

To explore whether mitochondrial function contributes to ALKBH1-dependent glioma phenotypes, we examined previously proposed links between ALKBH1 activity and mitochondrial metabolism^21,24,33,34^. Using Seahorse extracellular flux analysis, we assessed mitochondrial respiration in ALKBH1 KO and OE cell lines. Across all four models, genetic perturbation of ALKBH1 produced no or only minor changes in mitochondrial function (Figure 4a-b; Supplementary Figure 8), with the most notable effects being modest reductions in oxygen consumption in A172 and U251 KO cells (Supplementary Figure 8c and 8e). Consistent with these results, MitoTracker staining in transgenic 9L cells revealed no substantial differences in mitochondrial mass or distribution among KO, OE, and control cells (Figure 4c). The same was observed in the other cell lines examined (Data not shown). However, when mitochondrial function was challenged in galactose-containing media (a condition that forces reliance on oxidative phosphorylation rather than glycolysis), ALKBH1 KO cells exhibited the greatest sensitivity, indicating compromised OXPHOS capacity that is not apparent under standard culture conditions (Figure 4d). These findings suggest that while ALKBH1 does not dramatically alter baseline mitochondrial respiration or abundance, it contributes to maintaining mitochondrial fitness under metabolic stress, which may influence glioma cell survival and adaptability in vivo. Importantly, these effects cannot be attributed to mitochondrial f^5^C tRNA modifications, since ALKBH1 perturbations did not greatly affect their levels.

**Figure 4.**
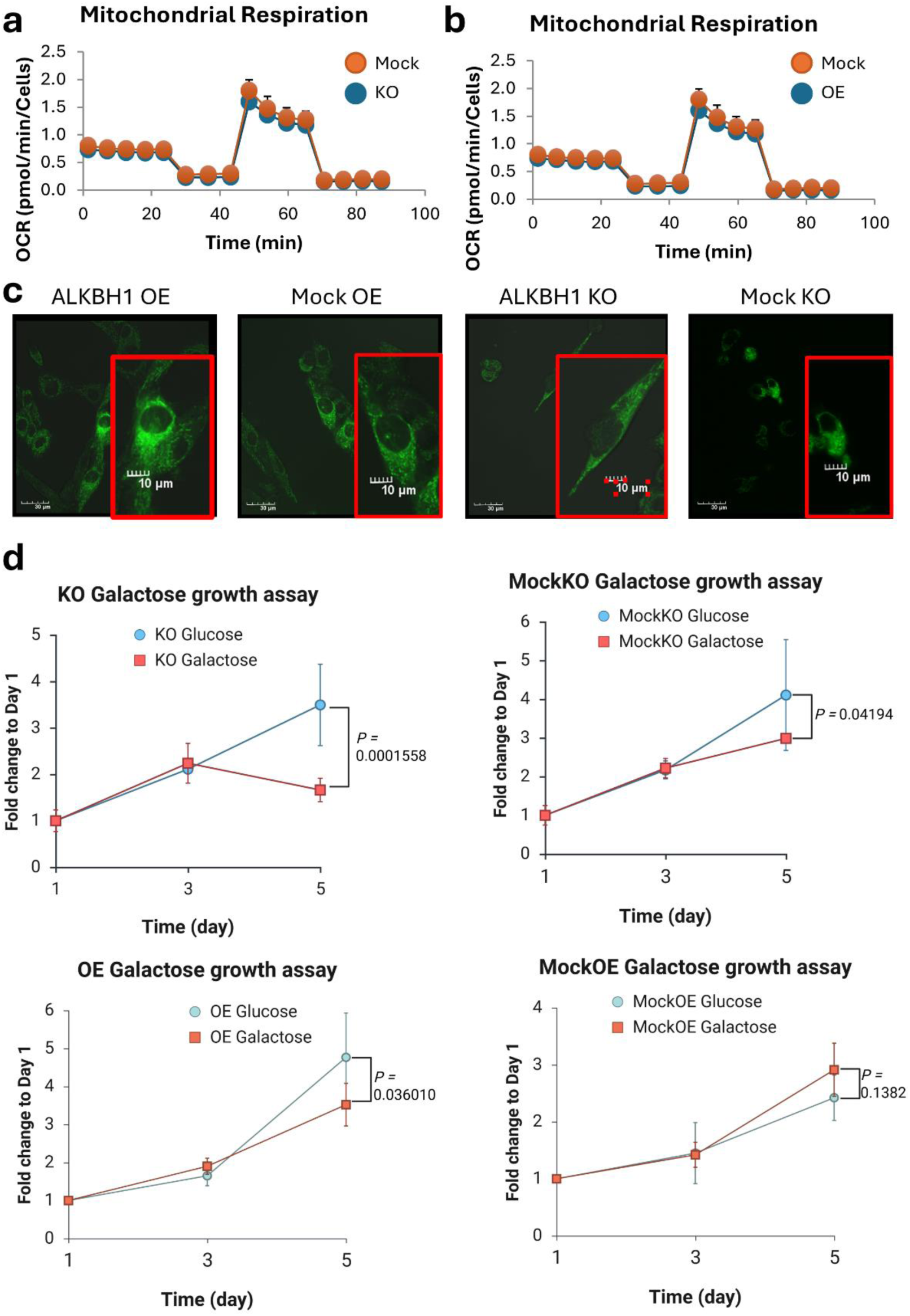
ALKBH1 is essential for mitochondrial fitness in glioma cells. **a, b**, Analysis of mitochondrial respiration using Seahorse flux analyzer in transgenic GS-9L cells. **c,** Mitotracker green live cell imaging in transgenic GS-9L cells. Bar size in photo = 30µm and in zoomed in inset = 10µm **d,** Analysis of transgenic GS-9L cell growth after galactose challenge. Linear regression was used for statistical analysis.

### ALKBH1 regulates codon decoding via anticodon modifications

To understand how ALKBH1 impacted glioma phenotype at the molecular level, we performed RNA-seq and Ribo-seq analysis on the 9L cell line used in the animal experiment. ALKBH1 KO led to minor transcriptional and translational changes, while OE had stronger transcriptional and translational impact (Figure 5a-b]. Spearman’s correlation analysis revealed no correlation between RNA-seq and Ribo-seq datasets in both KO and OE cells (Figure 6c-d]. Next, we focused on mRNA translational efficiency (TE) and analyzed the TE dataset (calculated by comparing the Ribo-seq to RNA-seq reads (see methods)) by pre-ranked gene set enrichment analysis (GSEA) of the gene ontology biological processes (GOBP) pathways. KO cells showed upregulation of pathways linked to neuronal and synaptic functions and downregulation of pathways linked to embryogenesis, while ALKBH1 overexpressing cells showed upregulation of pathways linked to genome replication and proliferation and downregulation of pathways linked to cholesterol and steroid biosynthesis (Supplementary Figure 9). These patterns could also indicate a shift towards stemness in OE cells while KO cells could be phenotypically shifted towards a more differentiated state, in line with our previous observations.

**Figure 5.**
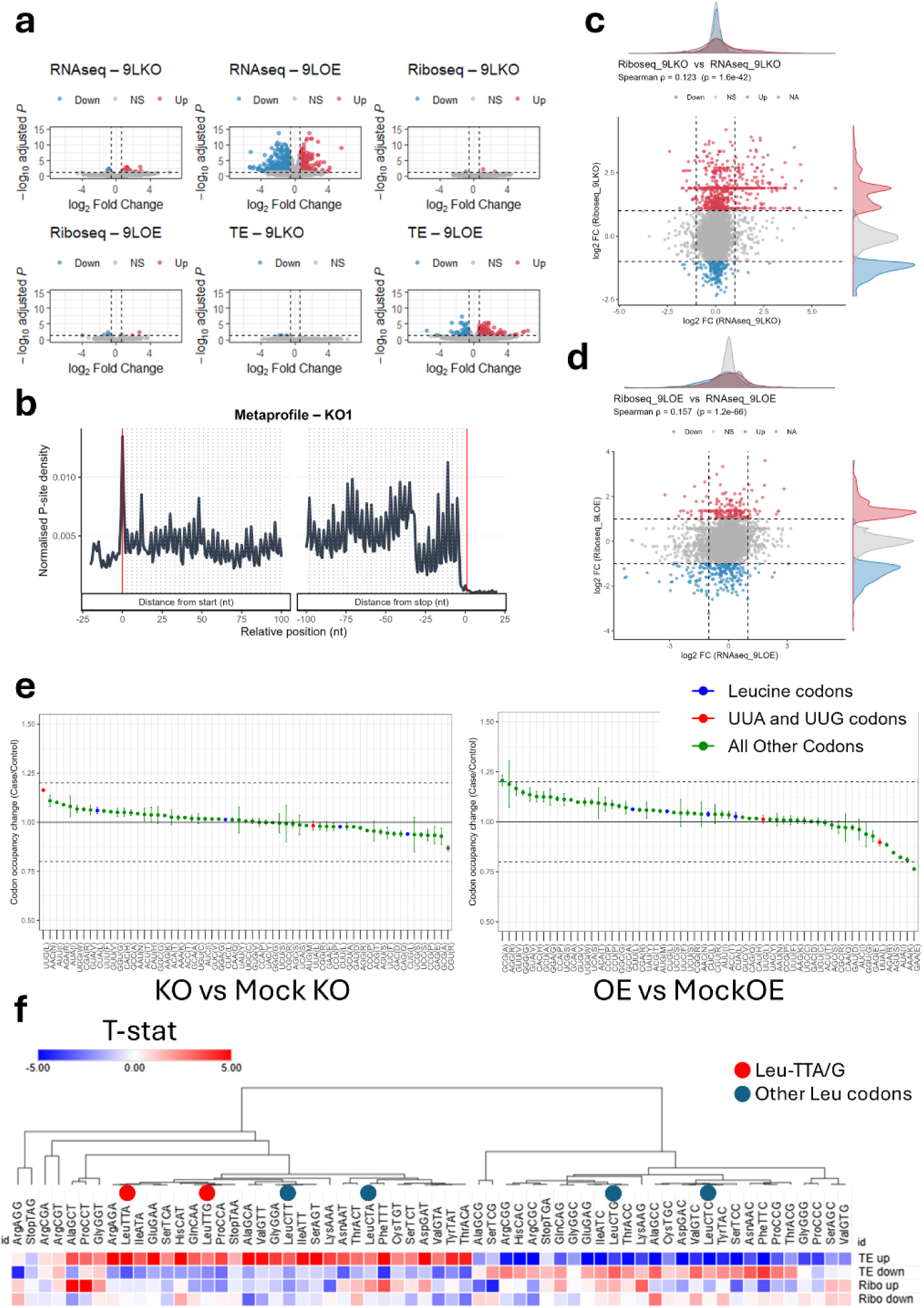
ALKBH1 promotes the translation of rare Leucine codons. **a**, Volcano plots showing the differentially expressed genes from RNA-seq, Ribo-seq, and translational efficiency (TE) analysis of GS-9L cells. Volcano plots show KO vs MockKO (9LKO plots) and OE vs MockOE (9lOE plots). **b,** Representative P-site metagene plot showing the 3-nucleotide periodicity observed in translating ribosomes in the Ribo-seq analysis. **c,** Spearman’s correlation analysis of log2FC genes values between RNA-seq and Ribo-seq dataset of KO cells. **d,** Spearman’s correlation analysis of log2FC genes values between RNA-seq and Ribo-seq dataset of OE cells. **e,** Ribosome A-site dwelling time (A-site pausing). **f,** Analysis of synonymous codon usage (isoacceptors frequencies) changes in OE vs MockOE datasets. The analysis focused on significantly differentially expressed genes in the Ribo-seq and TE datasets. Values represent T-stat, with |T-stat| ≥ 2 indicating statistically significant change in synonymous codon usage compared to the background (i.e. genome average).

**Figure 6.**
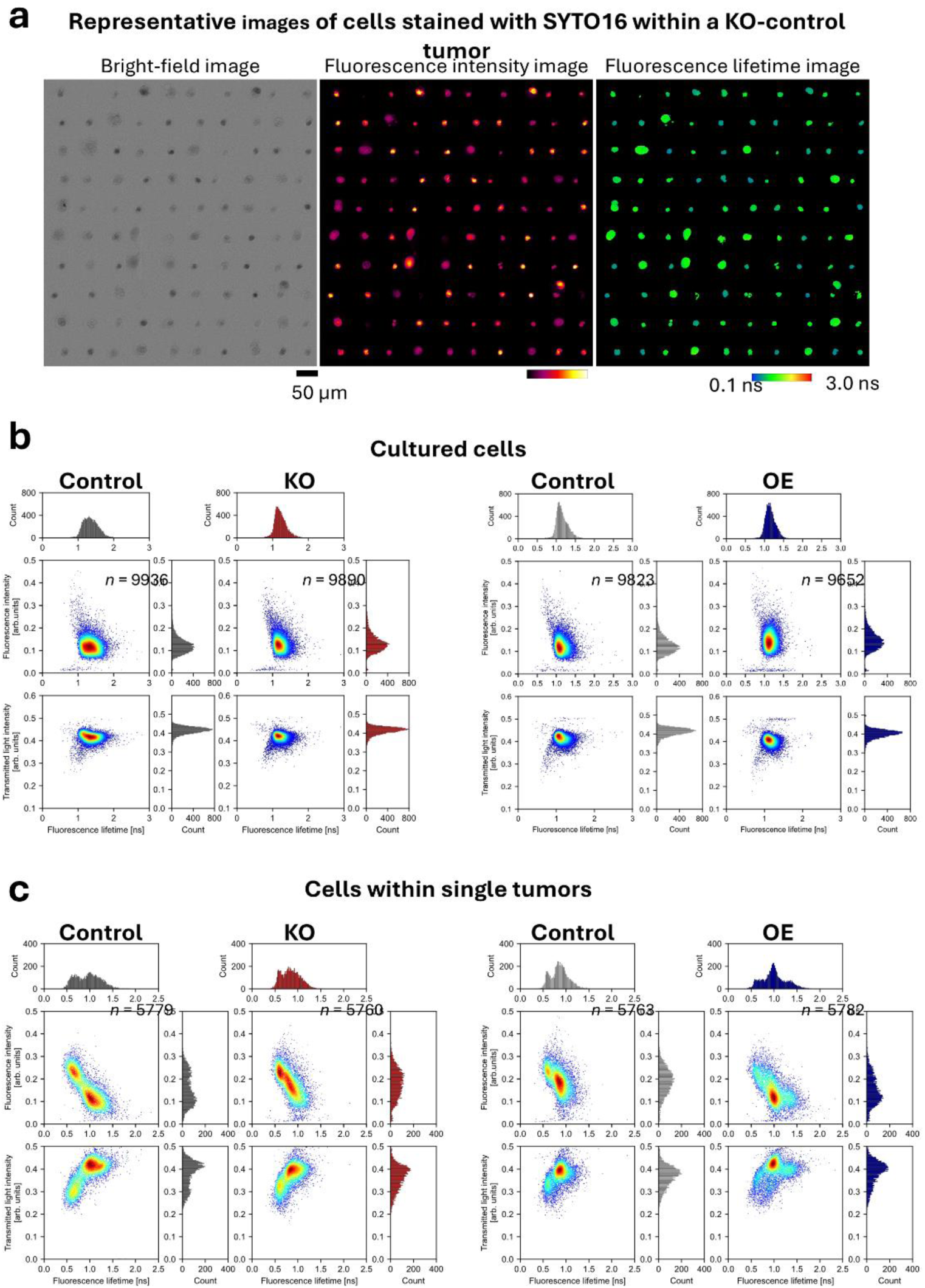
FLIM flow cytometry analysis. **a**, Representative bright-field, fluorescence, and fluorescence lifetime images of SYTO16-stained cells within a tumor (control for KO), acquired using a custom-built high-throughput FLIM flow cytometer. **b,** Distributions of cultured GS-9L cells (KO, OE, and their corresponding controls) stained with SYTO16, shown as scatter plots and histograms of transmitted light intensity, fluorescence intensity, and fluorescence lifetime derived from single-cell images. **c,** Distributions of intratumor cells (KO, OE, and their corresponding controls) stained with SYTO16, shown as scatter plots and histograms of transmitted light intensity, fluorescence intensity, and fluorescence lifetime derived from single-cell images.

Next, we interrogated the translational dynamics changes mediated by ALKBH1. Codon occupancy revealed changes in occupancy in KO cells and not OE cells in the form of increased occupancy across the coding sequences (CDS), especially downstream from the start codon (Supplementary Figure 10a). Additionally, we performed puromycin incorporation assay to analyze nascent protein synthesis to correlate it with the Ribo-seq analysis (Supplementary Figure 10b). OE cells showed significantly reduced nascent protein synthesis, in-line with their lower proliferation and shift towards stemness and self-maintenance. KO cells, on the other hand, showed a slight increase in protein synthesis, again correlating with the increased proliferation and with the increased elongating ribosome occupancy across the CDS.

Next, we examined ribosome dwelling times at the A-site, which are linked to codon-anticodon pairing dynamics. We observed some relative changes in ribosome decoding fidelity of Leu-UUA and Leu-UUG codons compared to other Leu codons (Figure 5e). In KO cells, we observed that Leu-UUG had the longest ribosome dwelling time of all codons. On the other hand, ALKBH1 OE rendered Leu-UUA the fastest translated codon of all Leu codons, followed by UUG. To further examine the impact of these changes on mRNA translation, we analyzed the differentially translated mRNAs for their codon sequences. We focused on OE datasets since KO did not lead to significant alterations in mRNA translation that passed our threshold (Fold change > 1.5 and FDR < 0.05). We examined the codon usage bias via three metrics: (i) GC3 scores, which indicates global third nucleotide codon bias^29^, (ii) isoacceptors codon frequencies which measure synonymous codon usage bias, and (iii) total codon frequencies, which compares the usage of any given codon to all codons in the coding sequence [See methods]. GC3 scores revealed a globally A/T-ending codon biased translational program after ALKBH1 overexpression (Supplementary Figure 11a) Importantly, this was only observed at the translation and translational efficiency levels and not at the RNA expression level. Next, we evaluated the codon bias using isoacceptors and total codon frequencies analysis to examine codon usage and optimality patterns with special attention to Leu-UUA and Leu-UUG codons that are decoded by f^5^C and hm^5^C. Leucine is decoded by 6 codons with variable frequencies of presence in the coding sequences. Importantly, Leu-UUA (Leu-TTG in the plots) is the rarest leucine codon. Isoacceptors frequencies analysis (i.e., synonymous codon usage analysis) revealed a shift in Leu codon optimality, with Leu-TTA being the most optimal codon, evident in the translational upregulation of mRNAs overusing it. Additionally, Leu-TTG, Leu-CTT, and Leu-CTA were also optimal, while the most abundant leucine codons in the genome, Leu-CTG and Leu-CTC were enriched in translationally silenced genes, indicating that the cells are underusing these codons (i.e., suboptimal codons) (Figure 5f). Total codon frequencies analysis showed the same pattern of leucine codons biases, with Leu-TTA, whose translation is dependent on ALKBH1 modifications system, to be the most optimal leucine codon (Supplementary Figure 11b). Thus, we can conclude that ALKBH1 overexpression drives a globally A/T-ending codon biased translational program, which was linked to oncogenic translational programs^29,35^, and shifts the codon optimality to prefer the rare Leu-TTA codons while rendering the abundant Leu-CTG suboptimal. Despite the global A/T-ending codon biased translation, Leu-TTG remained optimally translated, which indicates that hm^5^C and f^5^C modifications expand codon decoding of tRNA-Leu-CAA to Leu-TTA and maintain its interaction with its cognate Leu-TTG.

To understand why increased translation of Leu-TTA codon could support glioma oncogenesis, we analyzed rat coding sequences transcriptome-wide for their isoacceptors and total codon biases towards each of the leucine codons^29^. Isoacceptors and total codon frequencies analysis revealed similar distribution of Leu codons in the coding sequences across the rat genome (Supplementary Figure 12a and 13a). Next, we conducted GOBP overrepresentation analysis (ORA) for the top 5% genes based on their isoacceptors or total codon frequencies scores for each of the leucine codons. At the isoacceptors and total codon frequencies level analyses, we observed unique gene enrichment of different leucine synonymous codons, with some overlap between the two most used leucine codons in the genome, Leu-CTG and Leu-CTC (Supplementary Figure 12b and 13b). The same distribution was observed in human and mouse genomes as well (Data not shown). We further examined the enrichment of 3 of the leucine codons, the most used one in the genome (i.e., most optimal codon genomically), Leu-CTG, and the 2 codons decoded by hm^5^C/ f^5^C, Leu-TTG and Leu-TTA. At the isoacceptors level, we observed that genes overusing Leu-CTG functionally enrich for pathways related to differentiation and development as well as gas transport (Supplementary Figure 12c). Leu-TTG overusing genes enriched for pathways linked to RNA splicing and metabolism (Supplementary Figure 12d). Leu-TTA enriched genes functionally enriched for pathways linked to cell cycle and nuclear division (Supplementary Figure 12e). At the total codon frequencies level, Leu-CTG overusing genes were linked to membrane transport and G protein coupled receptor signaling. Leu-TTG overusing genes were linked to olfaction and taste responses. While Leu-TTA were linked to cell division and chromosome function linked pathways (Supplementary Figure 13c-e). Combined, our analysis reveals that a shift in Leucine codon optimality towards Leu-TTA followed by Leu-TTG leads to a pro-proliferative oncogenic program that maintains mitochondrial function and self-renewal capacity of cells, shifting glioma cells towards a more stem-like state and allowing the cells to fare better in the tumor microenvironment.

### ALKBH1 promotes glioma heterogeneity, stemness, immune evasion, and a neuronal phenotype

Our results pointed toward a prominent ALKBH1-driven glioma stem-like state. Given that we observed worse outcomes in the animal models without major changes in tumor sizes after Alkbh1 overexpression, we argued that tumor heterogeneity, a major contributor to glioma aggressiveness^4^, could be the cause of such worse outcomes. To test whether the expected changes in cellular heterogeneity mediated by ALKBH1 happen *in vitro* or *in vivo*, we analyzed our cell lines and tumors using High-throughput fluorescence lifetime imaging (FLIM) flow cytometry^36^. Here, we acquired large-scale images of whole cells using bright field, as well as of nuclei using fluorescence and fluorescence lifetime imaging, and analyzed these data to investigate how specific features emerge within the cell populations (Figure 6a). FLIM analysis revealed no changes in cell populations *in vitro* (Figure 6b). However, when examining tumors, we observed distinct subpopulations in ALKBH1 overexpressing cells derived tumors only, and not in other groups (Figure 6c).

Next, we used single cell RNA sequencing (scRNA-seq) to further assess and identify the cell populations withing the tumors. We sequenced glioma models generated using 9L cells (both KO and OE with their respective controls/Mocks). We were able to detect approximately 25,000 genes in 12,380 cells after filtering. Clustering revealed 17 distinct clusters (Figure 7a-b), which were annotated to 15 distinct cell types [Figure 7c] after identifying cluster specific markers (Supplementary Figure 14a). NG2 cells and oligodendrocyte precursor cells (OPCs) were the most abundant cell types detected followed by astrocytes and immune cells (tumor associated macrophages (TAMs), dendritic cells, macrophages, etc.) (Supplementary Figure 14b). We also detected several neuronal cell subtypes in the datasets such as ganglionic eminence cells, claustrum pyramidal cells and late GABAergic neurons and observed enrichment of several neuronal and synaptic markers such as Gja1 and Tenm2 (Figure 7d, Supplementary Figure 14b-d). We examined cell distribution across clusters to understand how ALKBH1 influences cell populations within the tumors. We argued that clusters that show enrichment in tumors generated by ALKBH1 overexpressing cells compared to their Mock and depletion or no change of ALKBH1 KO cells generated tumors compared to their Mock could indicate that ALKBH1 increases such cells in glioma tumors and vice versa (Figure 7e-f, Supplementary Figure 15a-b). For example, we observed that in the NG2 cells (neuron-glial antigen 2 expressing cells or polydendrocytes^37^) cluster, OE had more cells by percentage compared to MockOE, while KO had less cells than MockKO, indicating that ALKBH1 expression is linked to NG2 presence in the tumors. Collectively, ALKBH1 expression enhanced the presence of stem cells (Neural stem cells, NG2 cells, and oligodendrocytes precursor cells), neuronal cells (NG2 cells, spiral ganglion neurons, and Late GABAergic neurons), and tumor associated macrophages (TAMs), while it significantly reduced tumor infiltration by different types of tumor-suppressing immune cells such as NK T-cells and mature macrophages. Thus, in summary, ALKBH1 promotes a heterogenous, pro-neuronal, pro-glioma stem cells (GSCs), and pro-immune evasive phenotype in glioma tumors. We validated our observations using immunohistochemistry. We observed higher expression of the stemness markers Nestin, Sox2, and P53, and the neuronal/synaptic marker Ttyh1 in OE tumors and lower expression of Iba1 compared to other tumors (Supplementary Figure 16a-b). We also observed the downregulated of Nf (neurofibromatosis 1), a tumor suppressor^38^, in OE tumors compared to MockOE tumors (Supplementary Figure 16b). Collectively, this data confirms the increased stemness, aggressiveness, neuronal phenotype, and immune evasiveness brought about by ALKBH1 overexpression.

**Figure 7.**
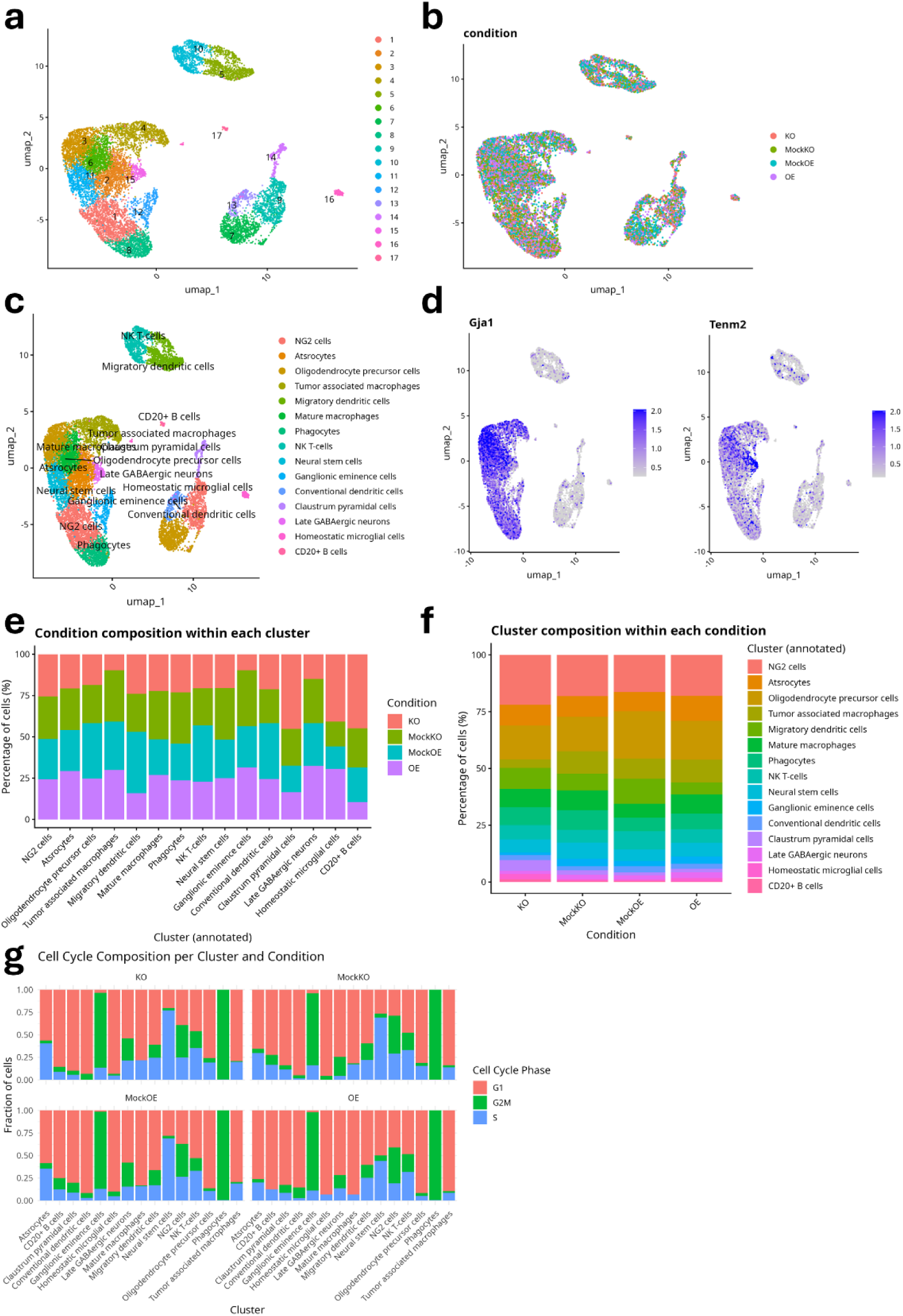
ALKBH1 alters tumor cell composition and microenvironment. **a**, UMAP clustering of scRNA-seq data showing the observed 17 cell clusters across the 4 groups. **b,** UMAP clustering showing the composition of different clusters by group. **c,** UMAP clustering with cluster cell annotations. **d,** Expression of Gja1 and Tenm2 across clusters. **e,** Ratio of cells belonging to each condition per cluster. **f,** Cluster composition within each condition. **g,** Cell cycle state across clusters and conditions.

**Figure 8.**
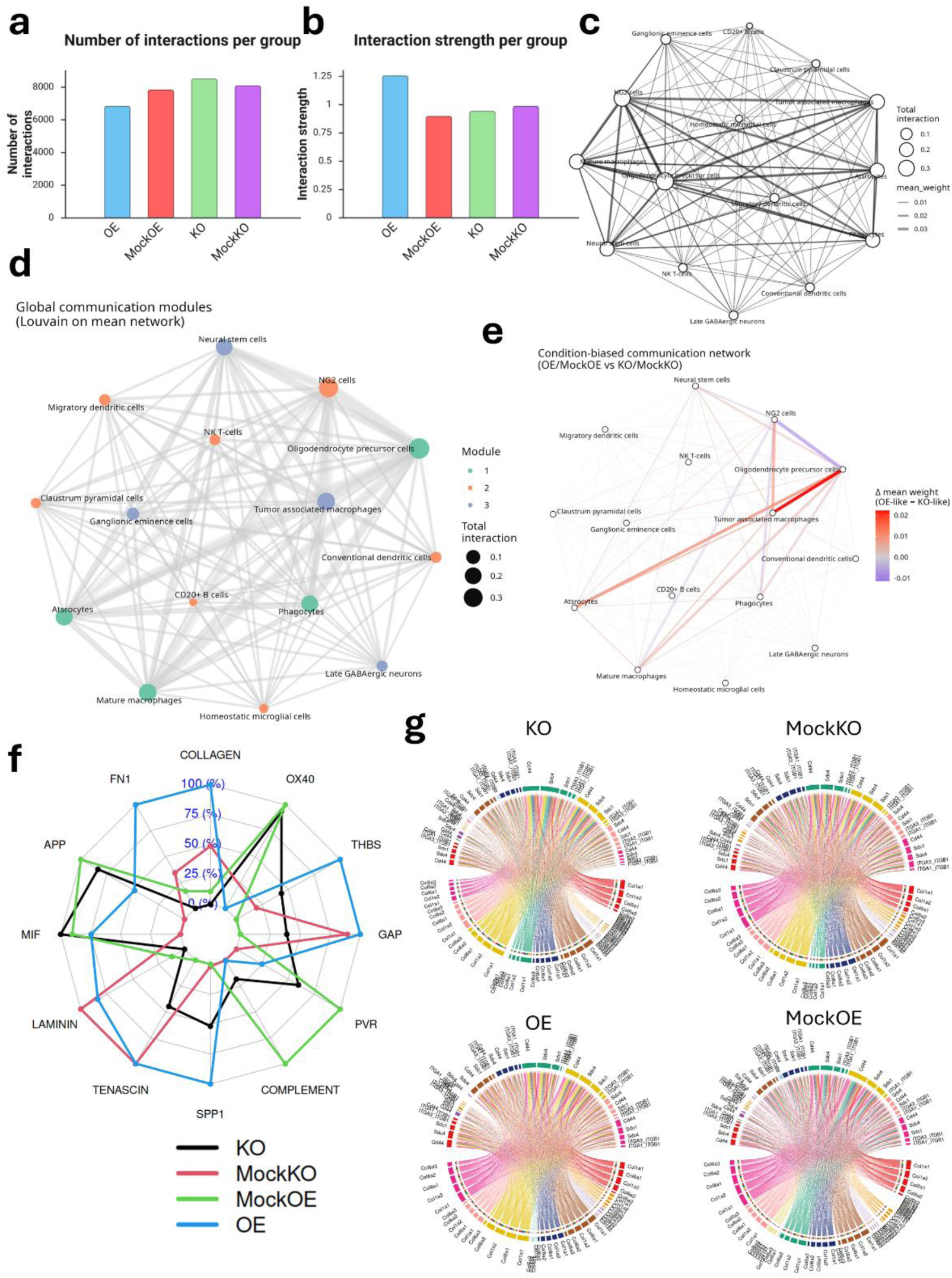
ALKBH1 regulates cell communication and glioma microenvironment. **a**, Number of interactions across different conditions. **b,** strength of interactions across different conditions. **c,** Global communication network strength across all conditions showing main communication hub cells. **d,** Louvain analysis on mean network revealed different modules of communications. **e,** Δ-network (OE vs KO bias). Condition biased communication network analysis (Red is stronger in OE/MockOE, Blue is stronger in KO/MockKO). **f,** Radar plot of top variable pathways across conditions showing the communication strength of each pathway. **g,** CellChat based analysis of COLLAGEN pathway interactions and strength in each condition. Cluster annotation in Supplementary Figure 18A.

To further understand the changes in the tumors beyond the cell proportions and distribution, we examined the cell cycle phases in each cluster (Figure 7g). We observed changes when comparing KO to MockKO and OE to MockOE tumors. For example, astrocyte cluster in KO tumors had higher proportion of cells in the S-phase compared to MockKO tumors, while OE tumors had higher proportion of cells in the G1-phase compared to MockOE. Neural stem cell cluster had more cells in the G1-phase in OE tumors compared to MockOE tumors. CD20+ B cell cluster had more cells in the G2M-phase in MockOE tumors compared to OE tumors where the cells were mostly in G1-phase and no G2M-phase cells could be detected. Overall, these observations reflect changes in cell states and phenotypes that are ultimately beneficial to the glioma cells, and that these changes are regulated by ALKBH1 either directly or indirectly via its role in regulating glioma cells mRNA translation.

Next, we wanted to understand the changes in cell state and tumor microenvironment as regulated by ALKBH1, we performed cell-to-cell communication analysis using CellChat^39^. We observed fewer but stronger interactions in the OE group compared to MockOE, while the KO group had more interactions compared to MockKO (Figure 8a-b). Detailed analysis of communication networks revealed evident changes in the numbers of interactions between distinct cell types. In KO vs MockKO comparison, we observed gain and loss of interactions between different cell types as a result of ALKBH1 KO (Supplementary Figure 17a), while in OE vs MockOE, we observed global reduction in the numbers of interactions, as expected, with selective activation of specific communication networks such as NG2 to Astrocytes and NG2 to TAMs (Supplementary Figure 17b). Global analysis of communication networks revealed that OPCs and NG2 cells act as main communication hubs, either as senders or receivers of signals (Figure 8b). It also showed extensive communication between OPCs and NG2 cells and mature macrophages and neural stem cells. Using Louvain method for community detection, we identified 3 modules of communications across different cell types (Figure 8d). OPCs, astrocytes, mature macrophages, and phagocytes formed one module, neural stem cells, TAMs, Late GABAergic neurons, and ganglionic eminence cells formed the second module, while the third was formed by the remaining cell types. Condition-biased network analysis revealed stronger communication between OPCs and TAMs, astrocytes, and mature macrophages in OE vs MockOE conditions indicating (Figure 8e).

Next, we examined the pathways regulating cell communication. Overall, we observed changes in pathway communication strengths between KO and OE tumors and their respective controls (Figure 8f, Supplementary Figure 17c). The major changes were observed in pathways linked to extracellular matrix (ECM) remodeling, followed by pathways related to immune cell activation. In OE tumors, there was extensive activation of ECM linked pathways, such as collagen, laminin, tenascin pathways and others, which were downregulated in KO cells (Supplementary Figure 17c). On the other hand, immune/inflammatory pathways, such as ICAM, VCAM, and others, were suppressed in OE tumors and activated in KO tumors, indicating increased immune activation and cytokine signaling (Supplementary Figure 17c). The collagen pathway, which was extensively activated in OE tumors (Figure 8f), showed the most extensive network in the OE cells, with the senders being astrocytes, TAMS, mature macrophages, and phagocytes, while the receivers being NG2 cells, OPCs, and neural stem cells (Figure 8g). On the other hand, KO tumors showed sparse collagen pathway network with fewer ligands. Similar patterns were observed in the Laminin signaling pathway (Supplementary Figure 18a), fibronectin pathway, and other pathways linked to ECM remodeling indicating a stiffer ECM in OE tumors^40^. Furthermore, the SPP1 pathway, which is linked to cancer stem cell maintenance and immune suppression^41^, was more activated in OE tumors (Supplementary Figure 18b). Overall, we observed that ALKBH1 overexpression reprograms the tumor microenvironment and cell communication networks towards a stem-like, stiff ECM-like, and immune-evasive state that supports glioma invasion, proliferation, and leads to worse outcomes.

Finally, to understand how changes in cell states could induce the changes in cell communication and phenotypes, we conducted differential gene expression analysis between KO and MockKO and OE and MockOE in each of the different cell clusters followed by ORA analysis. We focus on the more critical cell clusters that we observed to be master regulators of cell communication. OPCs showed a more robust differential gene expression changes in OE vs MockOE compared to KO vs MockKO (Supplementary Figure 19a-d). ORA analysis revealed that in KO vs MockKO the DEGs enriched for pathways mainly linked to cell cycle regulation. On the other hand, OE OPCs showed enrichment in pathways linked to ECM remodeling, cell adhesion, and immune interactions. Given that we identified OPCs as important hub for cell communication within the tumors, especially in OE tumors, such results confirm our observation of ALKBH1-driven ECM remodeling mediated by transcriptional changes in OPC cluster cells. We also observed changes in gene expression in immune cell clusters. NK T-cells, for example, showed no major differences between KO and MockKO conditions and no specific pathway enrichment (Supplementary Figure 19e), while in OE vs MockOE we observed downregulation in a number of genes that enriched for various NK related immune pathways, reflecting not only that ALKBH1 reduced NK T-cells infiltration, but also suppressed their activation and function (Supplementary Figure 19f-g). Together, these results reveal a unified mechanism through which ALKBH1 worsens glioma outcomes: by stabilizing a stem-like and neuronal-like tumor cell state, expanding intratumoral heterogeneity, stiffening and remodeling the ECM, disrupting immune infiltration and activation, and thereby fostering a microenvironment associated with highly aggressive glioma behavior^42,43^.

## DISCUSSION

Translational reprogramming is a fundamental oncogenic layer across cancers, enabling rapid proteome remodeling and phenotypic plasticity in response to stress and therapy pressure^9,44^. Multiple processes contribute to this mRNA translational reprogramming, including mRNA modifications, ribosome quality control, and, critically, tRNA expression and chemical modification^9,44^. In particular, tRNA-modification–driven codon-biased translation has emerged as a key mechanism whereby cells alter codon optimality to preferentially translate oncogenic transcripts that promote proliferation, invasion, immune evasion, and other malignant traits^9,11^. Despite the large catalog of human tRNA modifications, only a subset has been deeply characterized in cancer. Two well-studied modifications are 7-methylguanosine (m^7^G) at position 46, which enhances tRNA stability and supports oncogenic translation^9,18^, and N6-threonylcarbamoyladenosine (t^6^A) at position 37, recently shown to bolster glioma stem cell fitness and aggressiveness^15^. Consistent with these observations, we and others find that cancers, including glioblastoma, exhibit an A/T-ending codon-biased translational program (i.e., preference for codons with A or T at the third position)^29,35^. Both m^7^G (affecting many codons) and t^6^A (acting on ANN codons) can reinforce this bias^29^. However, most prior studies inferred modification activity indirectly from “writer” expression in cancer datasets, a strategy that misses key determinants such as tRNA abundance, modification stoichiometry, and metabolic inputs^11^ and therefore cannot accurately deduce the true modification state of tRNAs.

Motivated by these limitations, we took a tRNA-centric approach and directly quantified tRNA modifications in human gliomas by LC-MS/MS. This revealed broad enrichment of modifications in high-grade tumors irrespective of IDH status and confirmed several glioma-linked modifications (t^6^A, m^7^G, ms^2^i^6^A)^9,15,19^, while uncovering additional changes not previously studied in glioma or in other cancers. Importantly, expression of writer enzymes correlated poorly with modification levels, underscoring that genomic/transcriptomic approaches are insufficient to capture the epitranscriptomic landscape. Among the altered modifications, wobble-position modifications stood out as a plausible driver of codon-biased translation. We therefore focused on the ALKBH1–FTSJ1 pathway (Figure 1e), which converts wobble m^5^C to hm^5^C and f^5^C and then to hm^5^Cm, thereby expanding the decoding capacity of cytosolic tRNA-Leu-CAA from UUG to UUG/UUA and mitochondrial iMet-CAU from Met-AUG to Met-AUG plus Ile-AUA^21^. Nonetheless, our findings and analysis indicate that ALKBH1 predominantly acts via modulating leucine codons optimality in glioma with negligible impact on mitochondrial f^5^C status.

ALKBH1 is a dioxygenase originally implicated in DNA alkylation repair and histone oxidation, later recognized to modify tRNA. Its tRNA-centric function involves α-ketoglutarate- and O₂-dependent oxidation of wobble m^5^C to hm^5^C and f^5^C, which we validate here^21,45^. A second activity, namely demethylation of m^1^A at position 58, has been reported^22,24,25^; we did not observe m^1^A changes upon ALKBH1 loss across four glioma cell lines, consistent with the notion that this activity is context-specific or limited to mitochondrial tRNAs^21,24^. Prior claims that ALKBH1 demethylates DNA m^6^A are disputed by studies attributing signals to potential mycoplasma contamination or RNA misincorporation rather than genuine DNA modification^50–52^, and there is no compelling evidence that ALKBH1 directly demethylates mRNA (in contrast to ALKBH5/ALKBH3 acting on RNA^46^). Importantly, ALKBH1-linked wobble modifications have been connected to malignancy in other settings, including metastatic programs via mitochondrial f^5^C in head and neck cancer^47^ and leucine-codon optimality in acute myeloid leukemia^33^. In glioma, our data show that ALKBH1 reprograms translation at two levels: locally by boosting decoding of rare leucine codons (especially Leu-TTA) and globally by imposing an A/T-ending codon-biased program. This dual effect preferentially elevates translation of transcripts controlling cell-cycle progression, extracellular-matrix (ECM) remodeling, and immune suppression, features that align with enhanced stemness and aggressive tumor behavior.

Rather than focusing on a single downstream effector, our study reveals that ALKBH1 functions as a master regulator that rewires the translational bottleneck. By altering codon optimality, ALKBH1 initiates a cascade of cell-state transitions and microenvironmental changes. These include expansion of glioma stem-like populations, emergence of neuronal-like malignant states associated with poor prognosis, reinforcement of immune-evasive programs, and extensive ECM remodeling toward a stiff, pro-invasive niche. This multi-level reprogramming explains why ALKBH1 overexpression paradoxically slows proliferation in vitro yet markedly worsens survival in vivo; ALKBH1 shifts glioma cells towards a stem cell-like state that favors slower protein synthesis and self-maintenance. These findings position ALKBH1 as an attractive therapeutic target not only because of its cancer-specific enzymatic activity but also because its inhibition could collapse multiple oncogenic programs simultaneously by interrupting their shared translational foundation.

Despite these advances, several limitations warrant consideration. Our mechanistic analyses relied primarily on rodent and in vitro models, which do not fully recapitulate the spatial and immunological complexity of human gliomas. Although we confirmed ALKBH1-dependent hm^5^C/f^5^C formation, we were unable to validate ALKBH1-dependent demethylase activity on m^1^A, consistent with its proposed context-dependent behavior^22,24,25^. CellChat inference of ligand–receptor interactions highlights shifts in communication networks but requires further biochemical validation. Furthermore, our emphasis on global translational reprogramming means that specific downstream mRNA and protein targets remain to be mapped. Future studies integrating spatial transcriptomics, ribosome profiling of patient-derived glioma samples, CRISPR-based tRNA transcripts perturbations, and ALKBH1 enzymatic inhibitors will be crucial to establish further causal connections between ALKBH1 activity, codon bias, and glioma aggressiveness. Together, our findings reveal ALKBH1-mediated wobble oxidation as a potent driver of glioma progression and a promising therapeutic vulnerability in malignant gliomas.

## Methods

### Cell lines and culture conditions

GS-9L was purchased from ECACC (Cat# 94119705). U87MG (U87) was purchased from General Cell Collection (Cat# 89081402). A172 cell line was purchased from ECACC (Cat# 88062428). U251 cell line was purchased from ECACC (Cat# 09063001). Lenti-X 293T cells were obtained from Takara bio (Catalog #632180). GS-9L and A172 cell lines were cultured in Dulbecco’s modified Eagle’s medium (DMEM; Wako, Cat# 049-32645) supplemented 10% Fetal bovine serum (Thermofischer, Cat# A5256701), sodium pyruvate, sodium bicarbonate, and penicillin/streptomycin (P/S) (Nacalai Tesque, Cat# 09367-34). 293T cells were cultured in DMEM (Gibco, Cat# 10566016) supplemented with 10% FBS, P/S, and sodium pyruvate. U87 and U251 were cultured in E-MEM (Wako, Cat# 5508975) supplemented with 10% FBS and Penicillin/Streptomycin. Medium was replaced twice weekly, and cells were passaged once a week, at a maximum of 10 times from the point of recovery. All cells were incubated at 37℃ and 5% CO_2_. 293T cells were transfected for lentiviral production using Lipofectamine 3000 (Invitrogen, Cat# L3000015). The vector plasmids, pMD2.G (Addgene, Cat# 12259), and psPAX2 (Addgene, Cat# 12260) plasmids were triple transfected and incubated for 48 hours. After incubation, supernatants were used for downstream experiments.

### Generation of CRISPR KO cell lines

Single guide RNA sequences were synthesized and inserted into LentiCRISPR v2 plasmid (Addgene Cat# 52961). Target sequence against rat Alkbh1 was Forward: 5’CACCGGAGTACGGCGGACCTAGGAG and Reverse: 5’AAACCTCCTAGGTCCGCCGTACTCC.

Target sequence against human ALKBH1 was:

Forward: 5’CACCGGAAACCTAATGTATGTAACC

Reverse: 5’AAACGGTTACATACATTAGGTTTCC

After lentivirus production, the vector was transduced into cells and selected with puromycin. After selection, cells were seeded to 96 well plate with limiting dilution to obtain single cell clone. The knockout of target gene was verified using T7 endonuclease assay and western blotting.

### Generation of ALKBH1 overexpressing cell lines

Rat Alkbh1 (NM_001395611.1) or human ALKBH1 (NM_001395611.1) cDNA expressing lentiviral vectors or EGFP expressing vector for control were synthesized by VectorBuilder. After lentivirus production, vectors were transduced into cells and selection was performed using puromycin. After selection, protein overexpression was confirmed using western blotting.

### Proliferation analysis

Cell proliferation analysis was performed using WST- 8 assay (Nakalai tesque, Cat# 07553-44). Cells were seeded in 96 well culture plates at a density of 1 x 10^3^ cells per well in 100µL culture medium. At the designated time point, 10µL WST-8 were added to each well and the cells incubated for 4h at 37℃ and 5% CO2. The absorbance was measured at 450nm by Spectra Max plate reader (Molecular Devices Spectramax 190). The experiment was performed over 5 consecutive days, with cell proliferation measured on day 1, 3, and 5. Each group had 6 replicate wells (6 biological replicate per time point per group).

### Spheroid confirmation

All glioma cells cultured with normal medium were digested and seeded into 10 cm^2^ dishes at a concentration of 5 x 10^5^ cells/dish in 10mL stem cell culture medium. The stem cell culture medium was composed of NeuroCult Basal Medium (STEMCELL Technologies, Cat# ST-05700), NeuroCult Proliferation Supplement (STEM CELL Technologies, Cat# 05701), 20 ng/mL EGF (STEM CELL Technologies, Cat# 78006.1), 10 ng/mL bFGF (FUJIFILM Wako Pure Chemical Corporation, Cat# 06004543), and 2 µg/mL heparin (STEM CELL Technologies, Cat# 07980) without antibiotics. All cells were incubated at 37℃, and 5% CO^2^. 5 mL fresh stem cell culture medium was added every 3 days and cells were observed regularly under the microscope for spheroid formation for a duration of 7 days.

### Seahorse assay

Analysis of mitochondrial respiration was performed using Seahorse XF96 analyzer 96 (Agilent, CA, USA) and Seahorse XF Cell Mito Stress Test Kit (Agilent, Cat# 103015-100), Seahorse XFe96/XF Pro FluxPak (Agilent, Cat# 103792-100), and Seahorse XF DMEM medium (Agilent, Cat# 103575-100) to measure the oxygen consumption rate (OCR) and the extracellular acidification rate (ECAR). On the first day, we seeded 50,000 cells/well in 96 well cell culture plates and the XFe96 sensor cartridge was hydrated in 200μL calibration medium. On the next day, medium was replaced with Seahorse XF DMEM, supplemented with sodium pyruvate, L-Glutamine, and d-glucose. Cells were incubated at 37 °C in a CO ₂-free incubator for 1 hour before measurements. Over the measurement duration, cells were treated with preloaded mito-stress reagents, namely a final concentration of 1.5μM Oligomycin, 1μM carbonyl cyanide Carbonyl cyanide-4 (trifluoromethoxy) phenylhydrazone (FCCP), 0.5μM Antimycin A and Rotenone.

### Galactose assay

Cells were seeded at a density of 1,000 cells per well and cultured in glucose free DMEM (Wako, Cat# 042-32255) supplemented with 2mg/ml galactose. Viability was measured using WST-8 cell counting kit (Wako, Cat# 341-08001).

### Puromycin incorporation assay

After being incubated in a complete growth medium with 10 μg/ml puromycin (Sigma Aldrich, Cat# P8833) for 30 minutes, the cells were harvested as previously described and subjected to western blotting analysis using puromycin antibody.

### Patient samples collection

Glioma samples were obtained from excess surgical resection tissues in Tohoku University Hospital with written informed consent. Samples were trimmed and flash-frozen in liquid nitrogen immediately after extraction. The research was conducted according to the protocol that was approved by the ethical committee in Tohoku University (2023-1-446). All patient studies were conducted with the Declaration of Helsinki. The information of the collected samples used for tRNA modifications analysis are as follows:

- Astrocytoma, IDH-mutant, CNS WHO Grade 2, N=3.
- Astrocytoma, IDH-mutant, CNS WHO Grade 3, N=4.
- Astrocytoma, IDH-mutant, CNS WHO Grade 4, N=4.
- Oligodendroglioma, IDH-mutant, and 1p/19q-codeleted CNS WHO Grade 2, N=5.
- Oligodendroglioma, IDH-mutant, and 1p/19q-codeleted CNS WHO Grade 3, N=4.
- Glioblastoma, IDH-wildtype, CNS WHO Grade 4, N=11.

### Total RNA extraction and quality control

Cells were lysed using QIAzol Lysis Reagent (QIAGEN, Cat# 79306), and total RNA was extracted with the miRNeasy Mini Kit (QIAGEN, Cat# 217004) according to the manufacturer’s instructions. RNA purity and concentration were measured using a NanoDrop spectrophotometer (Thermo Fisher Scientific, Cat# ND-ONE-W), and RNA integrity was assessed using the RNA 6000 Nano Kit (Agilent, Cat# 5067-1511) on the Agilent 2100 Bioanalyzer. Only samples with a RNA Integrity Number (RIN) of 9 or higher were included in subsequent analyses.

### RNA sequencing

RNA sequencing (RNA-seq) was performed using two biological replicates per group, with mRNA enriched using the NEBNext Poly(A) mRNA Magnetic Isolation Module (NEB, Cat# E7490) and libraries prepared using the Ultra II Directional RNA Library Prep Kit (NEB, Cat# E7760) according to the manufacturer’s instructions. Library quality was assessed on an Agilent Bioanalyzer 2100 using the Agilent DNA 1000 kit (Cat# 5067-1504), and library concentrations were quantified with the NEBNext Library Quant Kit for Illumina (NEB, Cat# E7630); all RNA samples had RNA integrity numbers (RIN) greater than 7. Sequencing was conducted by Macrogen on an Illumina HiSeq X-ten platform using 150 bp paired-end reads. Raw Fastq files underwent quality control with FastQC, followed by trimming with Trimmomatic^48^ to remove adapter sequences and low-quality reads. Clean reads were aligned to the rattus norvegicus (Rn7) reference genome (UCSC) using the splice-aware aligner HISAT2^49^, and gene-level counts were generated with FeatureCounts^50^. Differentially expressed genes were identified using Limma-Voom with TMM normalization.

### Ribo-seq

Ribosome profiling was performed as previously described^51,52^. After sample collection, ribosome foot printing was carried out by adding 1.25 U/µL RNase I (NEB Cat# M0243L) to 500 µL of clarified lysate and incubating for 45 minutes at room temperature on a rotator. RNA was extracted using TRIzol reagent and the Qiagen miRNeasy kit, and ribosome-protected fragments (27–35 nt) were isolated by TBE-Urea gel electrophoresis. rRNA was depleted using the NEBNext rRNA Depletion Kit v2 (NEB Cat# E7400L), and end-repair of purified fragments was performed with T4 PNK followed by purification using Zymo Oligo Clean and Concentrator Kit (Zymo Research Cat# D4060). Sequencing libraries were prepared using the NEBNext Multiplex Small RNA Library Prep Kit for Illumina (NEB Cat# E7300S) and paired-end 150 bp sequencing was conducted on the Illumina HiSeq X Ten platform using two biological replicates per group. Ribo-seq data were processed as previously reported^53^, with adapter and low-quality read removal performed using SeqPrep and Trimmomatic. Reads were aligned first to rRNA and tRNA references (Rn6), then to the rattus norvegicus genome (Rn7) using Bowtie2. FeatureCounts was used to generate read count files, and differential expression analysis was conducted with Limma. Quality control was performed using RiboWaltz^54^. Codon occupancy and ribosome pausing were evaluated using RiboToolkit, and translational efficiency was calculated by normalizing Ribo-seq reads to RNA-seq reads using Riborex package (https://github.com/smithlabcode/riborex) with Limma-voom based approach. Pathway analysis and other downstream analyses were performed in R using appropriate packages.

### Codon analysis

Analysis of codon usage and optimality was performed as previously described^29,34,51,55^. All analysis was performed in R. First, we retrieved the rat CDS sequences from Ensemble. Next, we selected the longest transcript for each gene as the representative coding sequences and used the package coRdon to count the codons in each CDS to generate codon count tables. Next, we calculated the desired metrics using these formulae:

### GC3 score

The GC content at the third codon position (GC₃) was calculated per gene as:

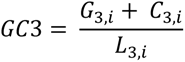

where 𝐺_3,𝑖_ + 𝐶_3,𝑖_ is the sum of codons having guanine or cytosine nucleotides at position 3 for gene

*i*, and 𝐿_3,𝑖_ the total number of codons in the gene.

### Isoacceptors codon frequencies

For each gene, codon usage was decomposed into frequencies of tRNA isoacceptors (synonymous codons decoding the same amino acid). Isoacceptors frequencies of a gene were calculated by:

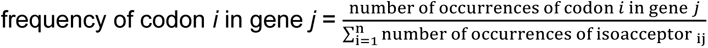

This yielded a score from 0 to 1.

### Total codon frequencies

Total codon frequencies compared the frequency of each codon to all other codons in a given sequence, which was calculated by:

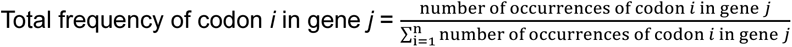

where n refers to the total number of codons used in the gene.

For the GC3 score analysis, up and downregulated genes were selected based on the desired thresholds (|log2FC| ≥ 1, FDR ≤ 0.05, relaxed FDR (in case fewer than 8 genes were significant) ≤ 0.1) and density plots were created to show the differences between the GC3 scores in the up or downregulated genes. In the case of isoacceptors frequencies or total codon frequencies, after selecting the up and downregulated genes, their frequencies were compared to the genome average using Welsh T-test to generate T-statistics (T-stat) values. |T-stat| ≥ 2 indicates statistical significance differences compared to the background (*p* < 0.05). The T-stat scores were used for creating heatmaps and visualization.

#### T-stat calculation

The t-statistics describing the codon frequency of a list of up- or down-regulated genes was calculated as:

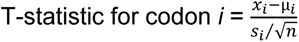

Where xi refers to the mean frequency for codon *i* in the sample, µ_𝑖_ refers to the mean frequency for codon *i* across the rattus norvegicus genome, si refers to the standard deviation of the frequency for codon *i* in the sample, and n refers to the sample size.

### Western blotting

Cells were homogenized in protein extraction reagent (Thermo Fisher Scientific, Cat# 78510), and the supernatant containing proteins separated. Protein concentrations were evaluated using Brachidonic acid assay kit (Thermo Fisher Scientific, Cat# 23227). Equal loads of proteins were separated on Mini-PROTEAN TGX gel (BIO-RAD, Cat# 4561035B05) and transferred to polyvinylidene difluoride membrane (BIO-RAD, Cat# 1704156) using semidry transfer. Membranes were blocked in iBind Flex solution kit (Thermo Fisher Scientific, Cat# SLF2020) and set iBind Flex Western Device (Thermo Fisher Scientific, Cat# SLF20002PK), following manufacturer protocol. The protein signal was detected using chemiluminescence Western Blotting detection reagents (Thermo Fisher Scientific, Cat# 32106) on ChemiDoc machine (BIO -RAD). Membranes were stripped and re-probed using anti-Actin-β antibody as a loading control. Antibodies were used anti-ALKBH1 antibody (1:1000 dilution, abcam, Cat# ab195376), anti-puromycin antibody (1:2000 dilution, Millipore, Cat# MABE343) anti-Sox2 antibody (1:1000 dilution, Cell Signaling technology, Cat# 2748S), anti-Oct4 antibody (1:5000 dilution, Proteintech, Cat# 60242-1-Ig), and anti-beta actin antibody (1:1000 dilution, Cell Signaling Technology, Cat# 4970S). The secondary antibody used was IgG detector solution, HRP-linked (1:400 dilution, Cell Signaling Technology, Cat# 7074S). The quantification of protein was measured by ImageJ software (National Institutes of Health, https://imagej.nih.gov/ij/).

### Liquid Chromatography mass spectrometry analysis of small RNA modifications

Analysis of small tRNA and small RNA modifications was performed as previously described^55^. Following small RNA (>90% tRNA) extraction from tissue or cell samples using purelink miRNA isolation kit (Thermo Fisher, Cat# K157001). Isolated small RNAs were digested in RNA digestion buffer (MgCl2 2.5mM, Tris 5mM (pH 8), Pentostatin 0.1μg/ml, deferoxamine 0.1mM, Butylated hydroxytoluene (BHT) 0.1mM, Benzonase 0.25 U/μl, Calf Intestinal Alkaline Phosphatase (CIAP) 0.1U/μl, and phosphodiesterase I (PDE I) 0.003 U/μl) for 6 hours at 37°C. After digestion, the digested RNAs were injected through a Waters BEH C18 column (50 × 2.1 mm, 1.7 µm) coupled to a Shimadzu Nextra Ultra-HPLC, with in-line UV detector, and a Shimadzu 8050 triple-quad mass spectrometer. The LC system was conducted at 25 °C and a flow rate of 0.35 mL/min. Buffer A was composed of 0.02% formic acid (FA) in DDW. Buffer B was composed of 70% Acetonitrile + 0.02% FA. Buffer gradient program was: 2min 0% B, 4min 5.7% B, 5.9min 72& B, 6min 100% B, 6.7min 100% B, 6.75min 0% B, 10min 0% B. The following source parameters were used: Nebulizing gas flow, 2.5 L/min, heating gas flow: 10 L/min, Interface temperature: 300°C, DL temperature: 150°C, heat block temperature: 400°C, Drying gas flow: 4 L/min, and interface voltage: 1kV. Loop time was set to 0.468sec, with a dwell time between 10 and 26 msec. The transition list can be found elsewhere^12^. The signals were normalized using the UV intensity of the canonical nucleosides to generate normalized peak areas for downstream analysis and a standard mix was used to determine the retention time and validate the results. All transitions were optimized beforehand for their collision energies and Q1 and Q3 biases. Data analysis was conducted using Lab solutions software.

### Mitotracker live cell staining

Live cell imaging was conducted with the MitoTracker™ Green FM (Invitrogen, Cat# M7514) using a glass bottom dish (Matsunami, Cat# D11131H). Confocal microscopy with Fluoroview v6 software (Olympus FV3000) was utilized to obtain images.

### Animal model of glioma

For intracranial allografts 10 weeks old adult male Fischer 344 rats weighing between 200 and 250g (Kumagai-Shigeyasu CO., Ltd) were randomly selected, and 50,000 glioma cells/2µL were stereotacticly implanted into the right striatum at a depth of 4.5mm following the standard procedure. Animals were carefully monitored after implantation until endpoint symptoms that determined previously were observed. Animals are housed in a controlled environment with a 12 h light/dark cycle, temperature regulated at 23℃ and fed in groups of 2 animals per cage. All animal experiments were conducted according to the procedures approved by the animal care facility of Tohoku University and according to the ARRIVE (Animal Research: Reporting In Vivo Experiments) guidelines. Ethical board approval was acquired before the commencement of this project (Approval number: 2024-Biomed-004-01).

### Tissue extraction and immunohistochemical analysis

Rats were euthanized by aspirating excess isoflurane 21 days after tumor cell implantation. They were first transcardially perfused with saline to remove the blood and subsequently with 2% paraformaldehyde in 0.1 mol/L saline. The fixed brain was removed and embedded in OCT compound (Tissue-Tek, Thermo Fisher Scientific, Cat# 4585) to be frozen by liquid nitrogen. The prepared brain was then sectioned at a 10μm thickness with a cryostat. Slides were created in coronal section, in which the tumor size was maximally shown. The sections were washed with 1 x PBS, incubated with blocking buffer with 20% Blocking Ace (UKB, Cat# UK880), 5% BSA (FUJIFILM Wako Pure Chemical Corporation, Cat# 010-25783), and 0.3% Triton X (Nacalai Tesque, Cat# 35501-15) in PBS for 1 h at room temperature before adding primary antibodies (details below) diluted in antibody-dilution buffer with 5% blocking Ace, 1% BSA and 0.3% Triton X in PBS, and incubated overnight at 4 °C. Subsequently, conjugated secondary antibodies of AlexaFluor 568 (1:500; Invitrogen, Cat# A11041, A10042, and A10037) and AlexaFluor 647 (1:500; Invitrogen, Cat# A21244, A21136, and 21244) were added and DAPI (1:500; Invitrogen, Cat# D1306) was used for counterstaining. Samples were inspected under a laser confocal microscope (OLYMPUS, FV3000).

Primary antibodies used:

- Anti-Ttyh1 antibody; 1:100, Proteintech, Cat# 26973-1-AP
- Anti-Nestin antibody; 1:100, Abcam, Cat# ab221660
- Anti-Iba1 antibody; 1:1000, Wako, Cat# 019-19541
- Anti-Sox2 antibody; 1:100, Abcam, Cat# 2748S
- Anti-P53 antibody; 1:50, Santa Cruz, Cat# sc-126
- Anti-Oct4 antibody; 1:250, Proteintech, Cat# 60242-1-Ig
- Anti-Nf antibody; 1:500, Abcam, Cat# ab72997
- Anti-MAP2; 1:5000, Abcam, Cat# ab5392

### Single-cell RNA sequencing

Rats were anesthetized by aspirating excess isoflurane 18 days after tumor implantation (GS-9L). They were transcardially perfused with saline to remove blood. Tumors were extracted from the rat brain. Tumor tissues were cut into pieces approximately 5mm in diameter and dissected into single cells by gentleMACS^TM^ Dissociator (Milteyi Biotec). Tissue cells were biotinylated using EZ-Link™ Sulfo-NHS-Biotin kit (Thermo Fisher Scientific Cat# 21217). The single cell suspensions were frozen and stocked with cell frozen buffer (CELLBANKER 1 plus, Takara Bio Cat# 11912). Samples were sent to ImmunoGeneTeqs (Chiba, Japan) and single-cell RNA sequencing was performed using the TAS-Seq platform. Libraries were sequenced on Illumina Novaseq-6000 platform using paired-end design that yielded a total of 705 million paired end reads. After trimming, quality filtering, and mapping to the Rattus norvegicus genome (rn6), 558 million paired end usable reads remained.

After demultiplexing and creation of read count matrices, 14,302 cells survived. The matrix was converted to a Seurat object for further downstream analysis.

### Single-cell RNA-seq Data Processing and Analysis

Single-cell RNA-seq data were processed in R (v4.3) using Seurat v5.0^56^. Cells with fewer than 200 or more than 7,000 detected genes and those with >5% mitochondrial transcripts and cell doublets were excluded. Gene expressions were normalized with LogNormalize (scale factor = 10,000). Data were scaled to zero mean and unit variance before principal component analysis (PCA). Batch correction and data integration across conditions (*KO*, *MockKO*, *MockOE*, *OE*) were performed using Harmony^57^ on the first 20 principal components (PCs). A shared nearest neighbor graph was constructed in the Harmony space, and clustering was carried out with the Leiden algorithm (resolution = 1.0). Two-dimensional UMAP embeddings were computed for visualization. Cluster-specific marker genes were identified using Seurat (Wilcoxon rank-sum test, *log2FC* > 1, *min.pct* > 0.25). Annotated clusters were assigned to canonical brain and immune cell types based on CellMarker 2.0 reference database^58^. Cluster proportions per condition were calculated, and differential enrichment was visualized with stacked bar plots. Cluster-specific differential gene expression (DEG) was conducted using Seurat’s FindMarkers function. Functional pathway enrichment of was performed using clusterProfiler v4.8. Cell-cycle phase scoring (S, G2/M) was performed with Seurat’s canonical markers.

Intercellular communication was inferred using CellChat v2.0^39^ with the CellChatDB.mouse ligand–receptor database. For each condition, communication probabilities and pathway-level strengths were computed following the standard CellChat manual. Multi-condition comparisons were generated with mergeCellChat(), producing rankNet, differential interaction, and pathway similarity UMAPs. Pathway-level strengths were aggregated across conditions into a pathway × condition matrix and visualized. Louvain community detection defined communication modules (glial–immune, myeloid–stromal, etc.), while a Δ-network contrasted *OE* vs *KO* conditions (edges colored by Δweight: red = *OE* enriched, blue = *KO* enriched) to reveal rewiring of cross-lineage signaling.

### FLIM flow cytometry analysis

Frozen transgenic GS-9L cell samples and frozen tumors from Fisher-344 rats using transgenic GS-9L cells were prepared in parallel with those used for single-cell RNA sequencing (stored separately) were transported to Tokyo University for FLIM flow cytometry analysis. After thawing, the cell samples were incubated in RPMI-1640 medium (FUJIFILM Wako Pure Chemical Corporation) supplemented with 10% FBS (MP Biomedicals) and 1% penicillin/streptomycin (FUJIFILM Wako Pure Chemical Corporation) at 37 °C in a 5% CO2 atmosphere. The incubation time was 0 min for cultured cells (Fig. 6B) or 7 h and 40 min for tumor-derived cells (Fig. 6A and 6C). After incubation, the cultured and tumor-derived cells were stained with 3 µM and 2.5 µM SYTO16 (Thermo Fisher Scientific), respectively, for 1 h, followed by FLIM flow cytometry analysis. Single-cell images were obtained using a custom-built high-throughput FLIM-based imaging flow cytometer, as described previously^36^. This system simultaneously captured bright-field, fluorescence intensity, and fluorescence lifetime images for each cell. The flow speed of cells during measurement was 1 m/s. The obtained images were analyzed using CellProfiler 4.2.1^59^, after aligning and concatenating multiple images in the same manner as shown in Fig. 6A. Nuclear regions were segmented from the fluorescence intensity images using a threshold set to 1/3.5 of the maximum intensity in each image, followed by morphological operations including dilation, erosion, and closing. For each nucleus, the mean transmitted light intensity, fluorescence intensity, and fluorescence lifetime were extracted.

### Statistical analysis and visualization

Statistical analysis was conducted using Graphpad Prism. ANOVA with Turkey’s post hoc, Student’s T-test, or linear regression were used depending on the data to be analyzed. All experiments were conducted with adequate number of biological replicates (stated in the results section or figure legends). Visualization was performed in *R* language (RStudio) or using BioRender.

## Acknowledgments

This work was funded by the Japan society for promotion of science (JSPS) grants numbers 23H02741 and 20KK0338 for Rashad, number 24K18149 for Kanno, and number 25K21826 for Niizuma. The work was also funded by JST Moonshot R&D project number JPMJPS2023 for Niizuma and Takeda Science Foundation grant for Kanno. We thank Serendipity Lab for providing collaboration opportunities. The authors would like to thank Natsumi Konno and Michitoshi Watanabe for aiding with this work.

## Ethics declarations

K.G. is a shareholder of CYBO, LucasLand, and FlyWorks.

## Data availability

Raw and aligned sequencing data were deposited in the sequence read archive (SRA) under the accession number: PRJNA1358817

## Author information

Contributions

S.R. conceived and designed this study. A.N., A.N., Y.I., A.M., M.K., M.K., S.A.-M., and H.K. conducted the experiments. S.R. and A.N. performed bioinformatics and sequencing data analysis. H.K. conducted the FLIM experiment and data analysis. A.N., M.K., S.Y., Y.S., H.E., and K.N. obtained the glioma clinical data and collected the samples. H.K., K.G., S.R., and K.N. obtained funding. S.R. drafted the manuscript. D.A., K.N., M.K., H.E., and K.G. critically revised the manuscript. All authors approved the final version of the manuscript.

## Inclusion and ethics

This study followed ethical guidelines with informed consent obtained for all samples and protocols approved by institutional ethics committees. Data was analyzed with awareness of potential biases. We are committed to promoting equity and inclusion in research while advancing scientific understanding.

