## Supplementary figures and tables for "ALKBH1 drives codon-biased, pro-oncogenic translation and tumor microenvironment remodeling in glioma via tRNA wobble oxidation": Supplementary figures.pdf

**Supplementary figure 1: Analysis of the correlation between ALKBH1 and FTSJ1 expression and glioma outcomes in 2 databases, UCSC Xena (https://xena.ucsc.edu/welcome-to-ucsc-xena/) and the human protein atlas (https://www.proteinatlas.org/).**. **a:** Kaplan-Meier curve showing the correlation between ALKBH1 expression and glioma outcomes. Data from the human protein atlas data. **b:** Kaplan-Meier curve showing the correlation between ALKBH1 expression and all glioma outcomes. Data from UCSC Xena. **c:** Kaplan-Meier curve showing the correlation between ALKBH1 expression and glioblastoma outcomes. Data from UCSC Xena. **d:** Kaplan-Meier curve showing the correlation between ALKBH1 expression and low-grade glioma (LGG) outcomes. Data from UCSC Xena. **E-h:** the same analysis done in **a-d** repeated for JTSJ1. P values represent results from log-rank test.

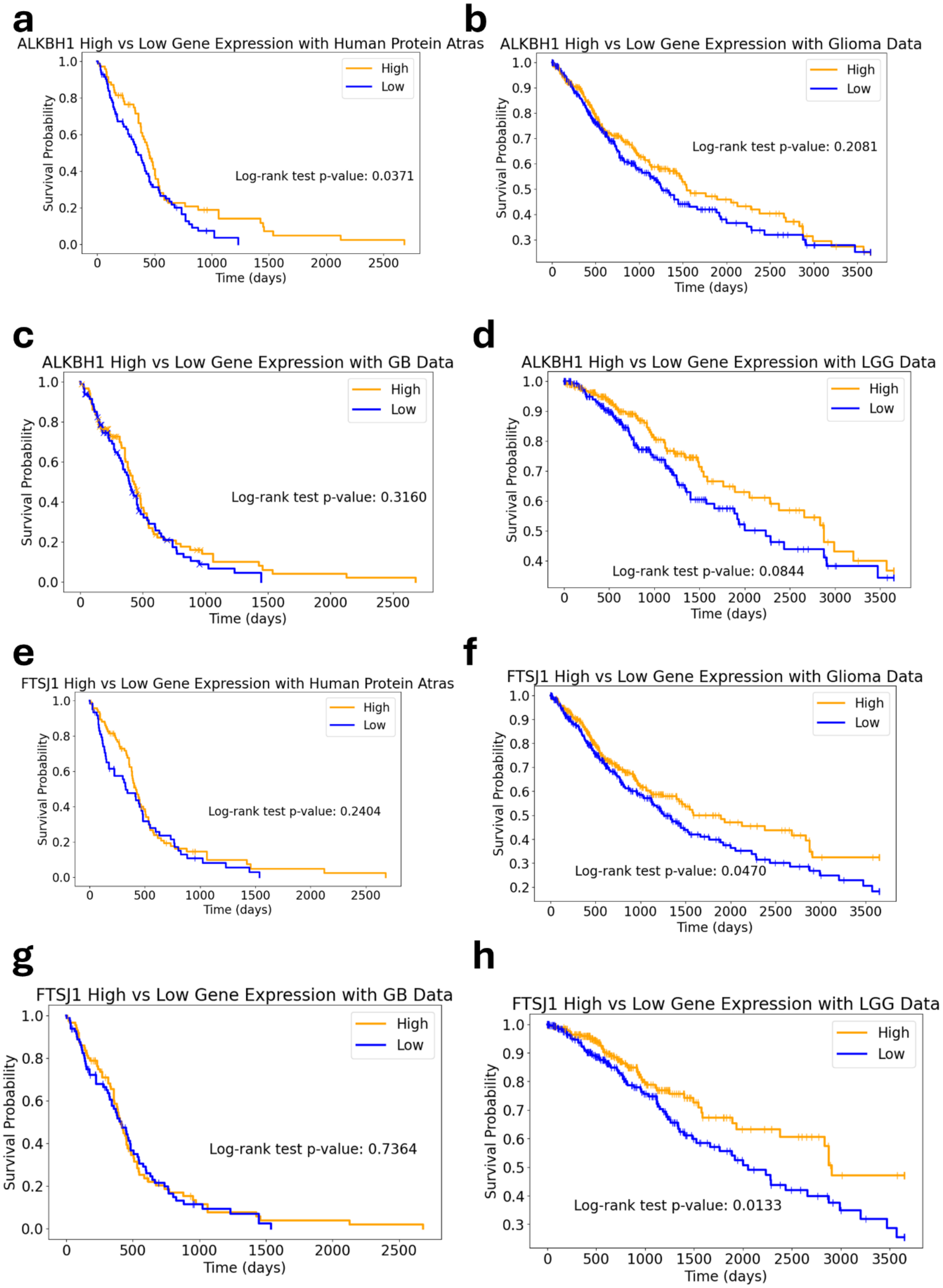
a
ALKBH1 High vs Low Gene Expression with Human Protein Atras
High
Low
Log-rank test p-value: 0.0371
Survival Probability
Time (days)
b
ALKBH1 High vs Low Gene Expression with Glioma Data
High
Low
Log-rank test p-value: 0.2081
Survival Probability
Time (days)
c
ALKBH1 High vs Low Gene Expression with GB Data
High
Low
Log-rank test p-value: 0.3160
Survival Probability
Time (days)
d
ALKBH1 High vs Low Gene Expression with LGG Data
High
Low
Log-rank test p-value: 0.0844
Survival Probability
Time (days)
e
FTSJ1 High vs Low Gene Expression with Human Protein Atras
High
Low
Log-rank test p-value: 0.2404
Survival Probability
Time (days)
f
FTSJ1 High vs Low Gene Expression with Glioma Data
High
Low
Log-rank test p-value: 0.0470
Survival Probability
Time (days)
g
FTSJ1 High vs Low Gene Expression with GB Data
High
Low
Log-rank test p-value: 0.7364
Survival Probability
Time (days)
h
FTSJ1 High vs Low Gene Expression with LGG Data
High
Low
Log-rank test p-value: 0.0133
Survival Probability
Time (days)

**Supplementary figure 2:** Western blotting validation of ALKBH1 expression after KO or OE in U87, U251, and A172 cells. Graphs show the changes in expression as fold change to appropriate Mock. Student's t-test was used for statistical analysis (N = 3 biological replicates per group).

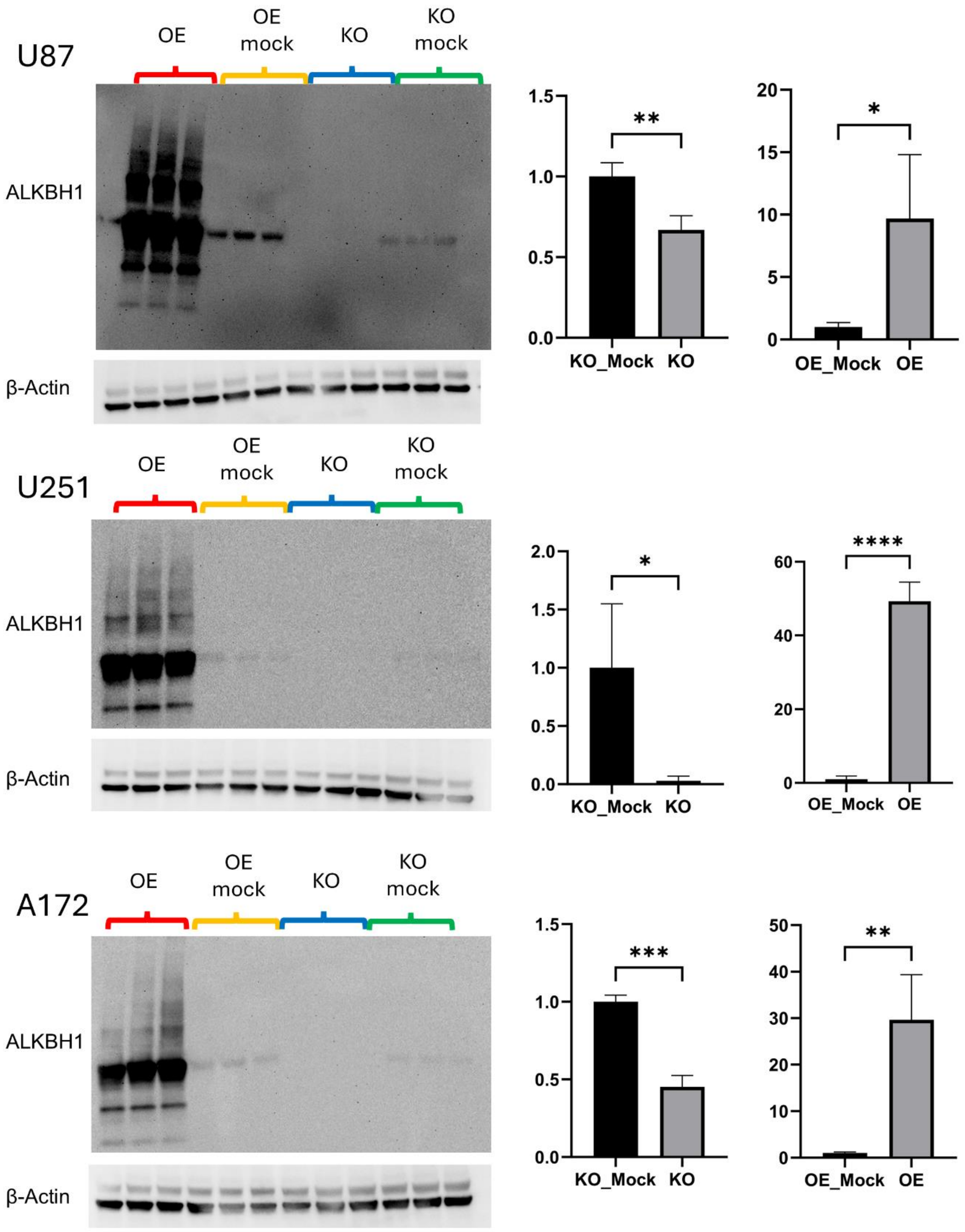
U87
OE
OE mock
KO
KO mock
ALKBH1
β-Actin
1.5
1.0
0.5
0.0
**
KO_Mock
KO
20
15
10
5
0
*
OE_Mock
OE
U251
OE
OE mock
KO
KO mock
ALKBH1
β-Actin
2.0
1.5
1.0
0.5
0.0
*
KO_Mock
KO
60
40
20
0
****
OE_Mock
OE
A172
OE
OE mock
KO
KO mock
ALKBH1
β-Actin
1.5
1.0
0.5
0.0
***
KO_Mock
KO
50
40
30
20
10
0
**
OE_Mock
OE

**Supplementary figure 3:** Detailed graphs of the changes in ALKBH1-related modifications in GS-9L cells. **a:** KO cells graphs. **b:** OE cells graphs. Statistical analysis: unpaired t-test. *: < 0.05. ** < 0.005. *** < 0.0005. **** < 0.0001. N = 3 biological replicates. Y axis: Normalized Peak areas.

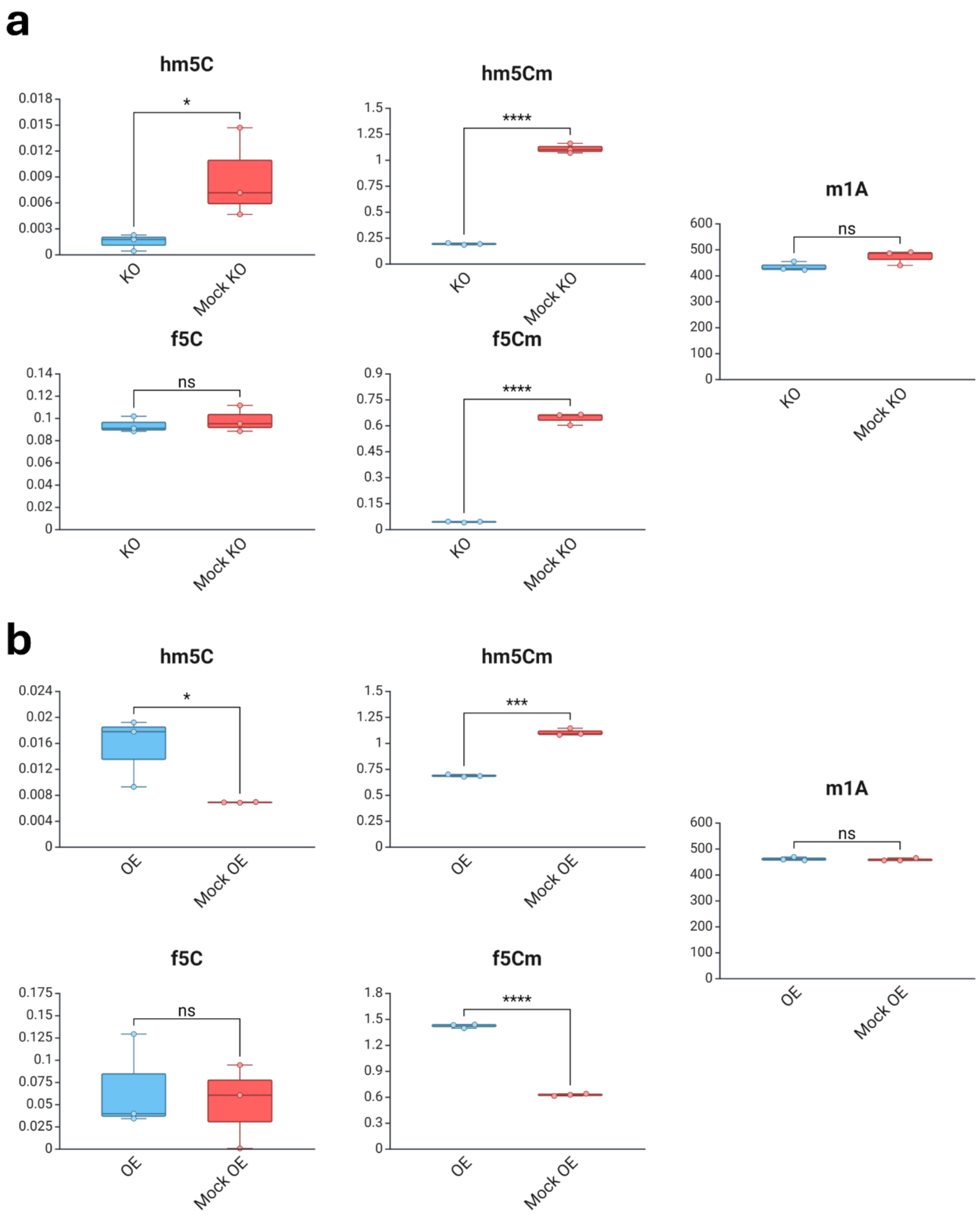
a
hm5C
0.018
0.015
0.012
0.009
0.006
0.003
0
*
KO
Mock KO
hm5Cm
1.5
1.25
1
0.75
0.5
0.25
0
****
KO
Mock KO
m1A
600
500
400
300
200
100
0
ns
KO
Mock KO
f5C
0.14
0.12
0.1
0.08
0.06
0.04
0.02
0
ns
KO
Mock KO
f5Cm
0.9
0.75
0.6
0.45
0.3
0.15
0
****
KO
Mock KO
b
hm5C
0.024
0.02
0.016
0.012
0.008
0.004
0
*
OE
Mock OE
hm5Cm
1.5
1.25
1
0.75
0.5
0.25
0
***
OE
Mock OE
m1A
600
500
400
300
200
100
0
ns
OE
Mock OE
f5C
0.175
0.15
0.125
0.1
0.075
0.05
0.025
0
ns
OE
Mock OE
f5Cm
1.8
1.5
1.2
0.9
0.6
0.3
0
****
OE
Mock OE

**Supplementary figure 4:** Detailed graphs of the changes in ALKBH1-related modifications in U87 cells. **a:** KO cells graphs. **b:** OE cells graphs. Statistical analysis: unpaired t-test. *: < 0.05. ** < 0.005. *** < 0.0005. **** < 0.0001. N = 3 biological replicates. Y axis: Normalized Peak areas.

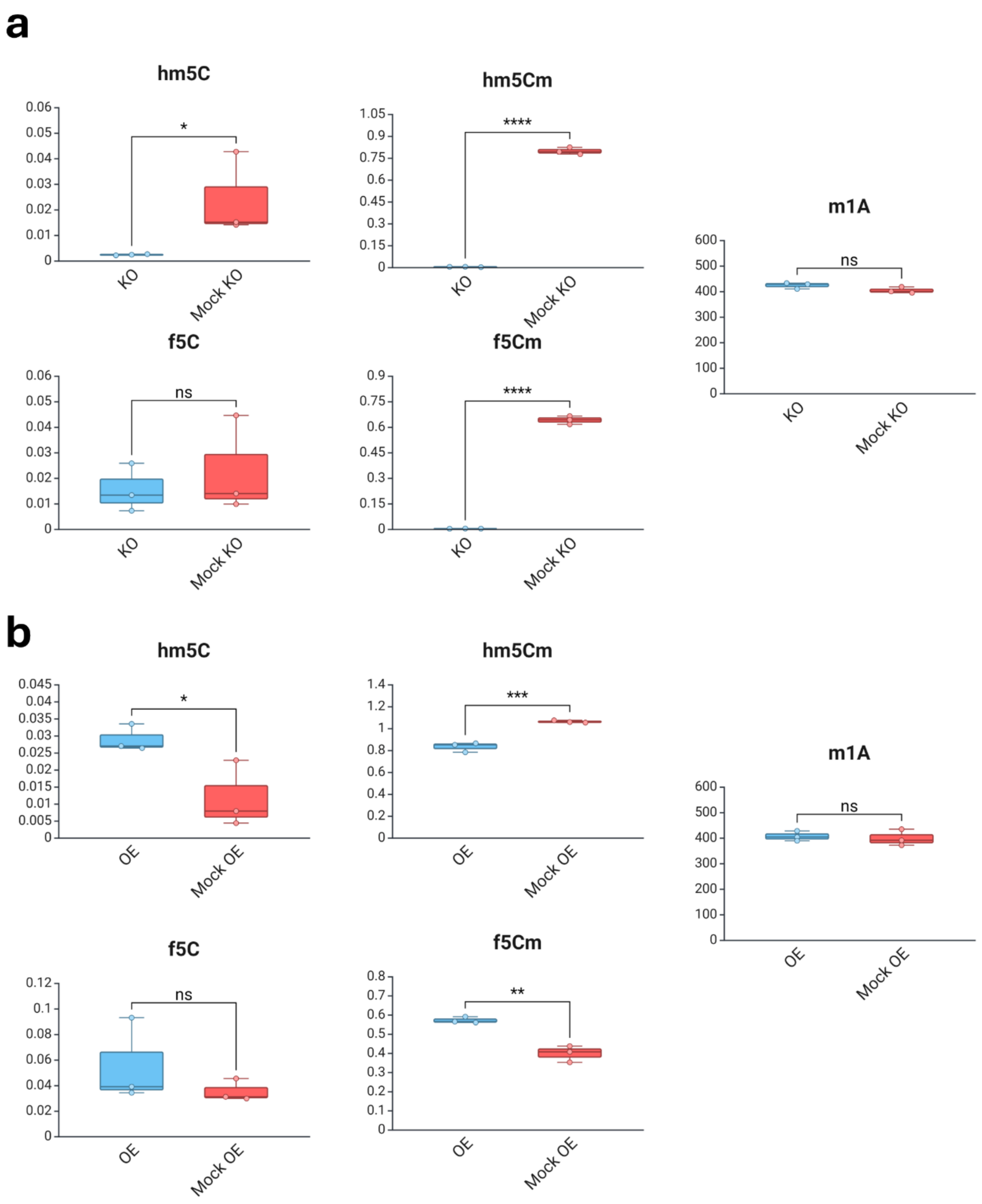

a
hm5C
hm5Cm
m1A
f5C
f5Cm
KO
Mock KO
*
****
ns
b
hm5C
hm5Cm
m1A
f5C
f5Cm
OE
Mock OE
*
***
**
ns

**Supplementary figure 5:** Detailed graphs of the changes in ALKBH1-related modifications in A172 cells. **a:** KO cells graphs. **b:** OE cells graphs. Statistical analysis: unpaired t-test. *: < 0.05. ** < 0.005. *** < 0.0005. **** < 0.0001. N = 3 biological replicates. Y axis: Normalized Peak areas.

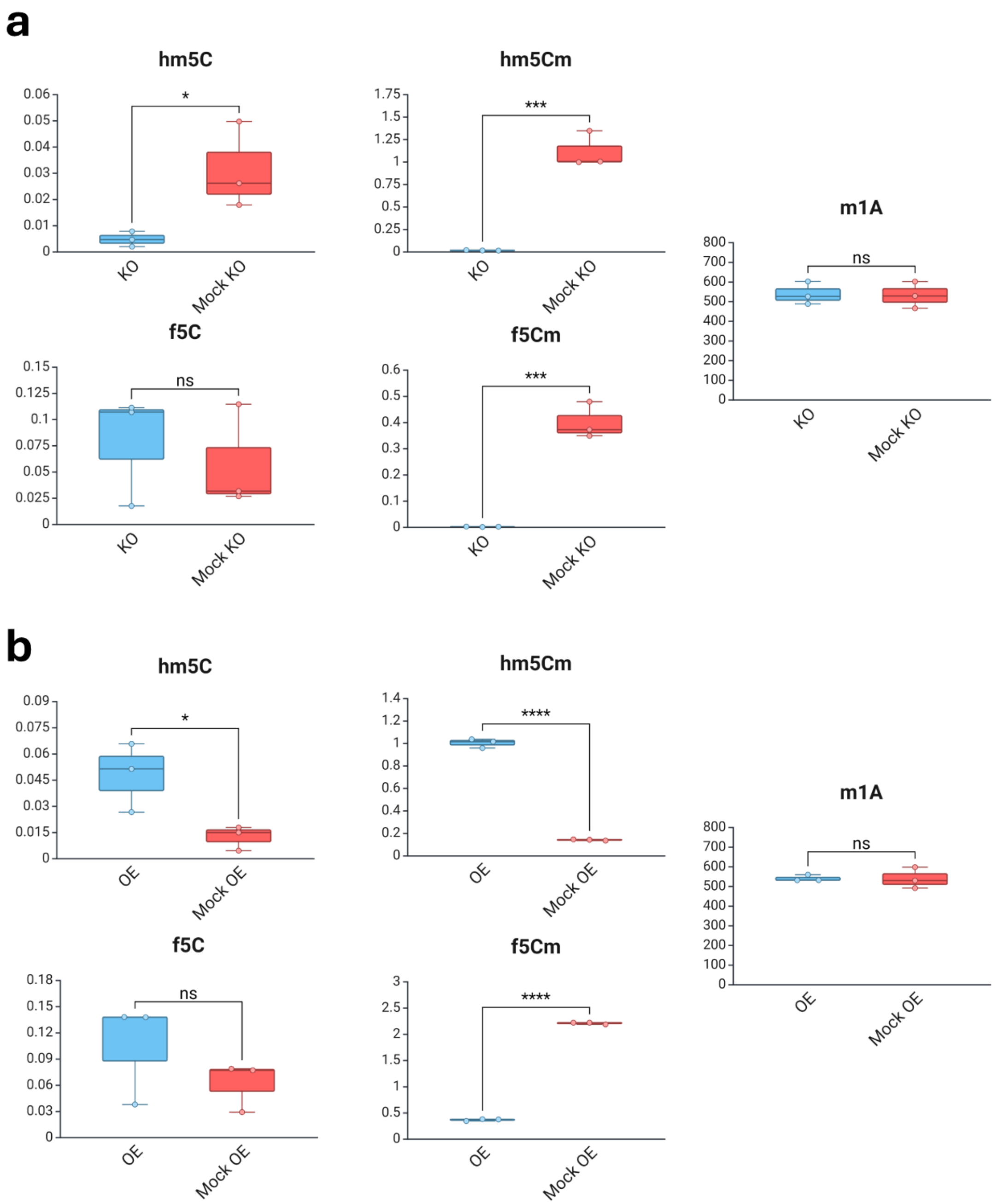

a
hm5C
*
KO
Mock KO
hm5Cm
***
KO
Mock KO
m1A
ns
KO
Mock KO
f5C
ns
KO
Mock KO
f5Cm
***
KO
Mock KO
b
hm5C
*
OE
Mock OE
hm5Cm
****
OE
Mock OE
m1A
ns
OE
Mock OE
f5C
ns
OE
Mock OE
f5Cm
****
OE
Mock OE

**Supplementary figure 6:** Detailed graphs of the changes in ALKBH1-related modifications in U251 cells. **a:** KO cells graphs. **b:** OE cells graphs. Statistical analysis: unpaired t-test. *: < 0.05. ** < 0.005. *** < 0.0005. **** < 0.0001. N = 3 biological replicates. Y axis: Normalized Peak areas.

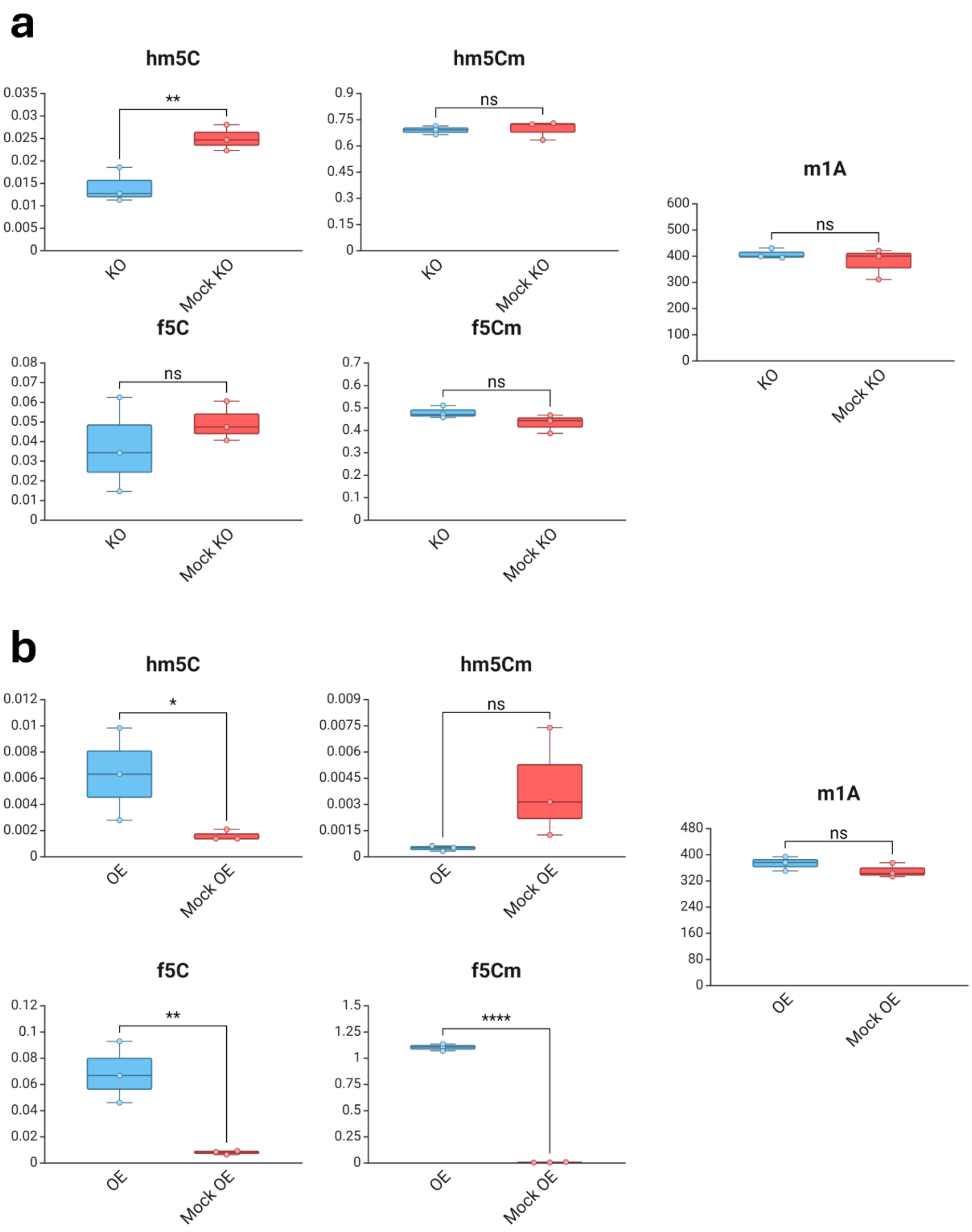
a
hm5C
0.035
0.03
0.025
0.02
0.015
0.01
0.005
0
**
KO
Mock KO
hm5Cm
0.9
0.75
0.6
0.45
0.3
0.15
0
ns
KO
Mock KO
m1A
600
500
400
300
200
100
0
ns
KO
Mock KO
f5C
0.08
0.07
0.06
0.05
0.04
0.03
0.02
0.01
0
ns
KO
Mock KO
f5Cm
0.7
0.6
0.5
0.4
0.3
0.2
0.1
0
ns
KO
Mock KO
b
hm5C
0.012
0.01
0.008
0.006
0.004
0.002
0
*
OE
Mock OE
hm5Cm
0.009
0.0075
0.006
0.0045
0.003
0.0015
0
ns
OE
Mock OE
m1A
480
400
320
240
160
80
0
ns
OE
Mock OE
f5C
0.12
0.1
0.08
0.06
0.04
0.02
0
**
OE
Mock OE
f5Cm
1.5
1.25
1
0.75
0.5
0.25
0
****
OE
Mock OE

**Supplementary figure 7: a-f:** Analysis of proliferation after ALKBH1 KO or OE in U87 (**a-b**), U251 (**c-d**), and A172 (**e-f**) cell lines. Linear regression was used for statistical analysis.

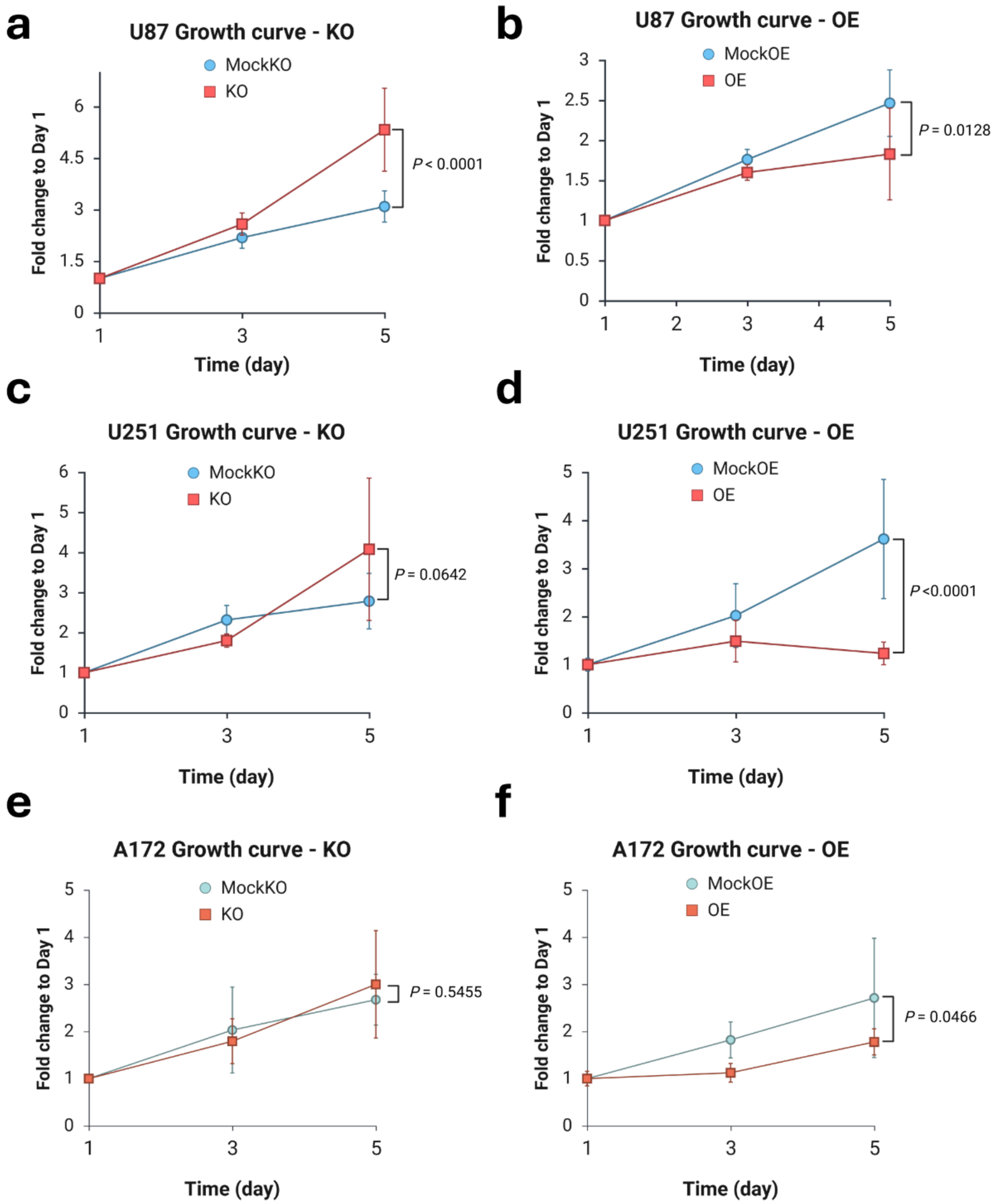
a
U87 Growth curve - KO
MockKO
KO
P < 0.0001
Fold change to Day 1
Time (day)
b
U87 Growth curve - OE
MockOE
OE
P = 0.0128
Fold change to Day 1
Time (day)
c
U251 Growth curve - KO
MockKO
KO
P = 0.0642
Fold change to Day 1
Time (day)
d
U251 Growth curve - OE
MockOE
OE
P <0.0001
Fold change to Day 1
Time (day)
e
A172 Growth curve - KO
MockKO
KO
P = 0.5455
Fold change to Day 1
Time (day)
f
A172 Growth curve - OE
MockOE
OE
P = 0.0466
Fold change to Day 1
Time (day)

**Supplementary figure 8:** Analysis of mitochondrial respiration using Seahorse flux analyzer in transgenic U87 (**a-b**), A172 (**c-d**), and U251 (**e-f**).

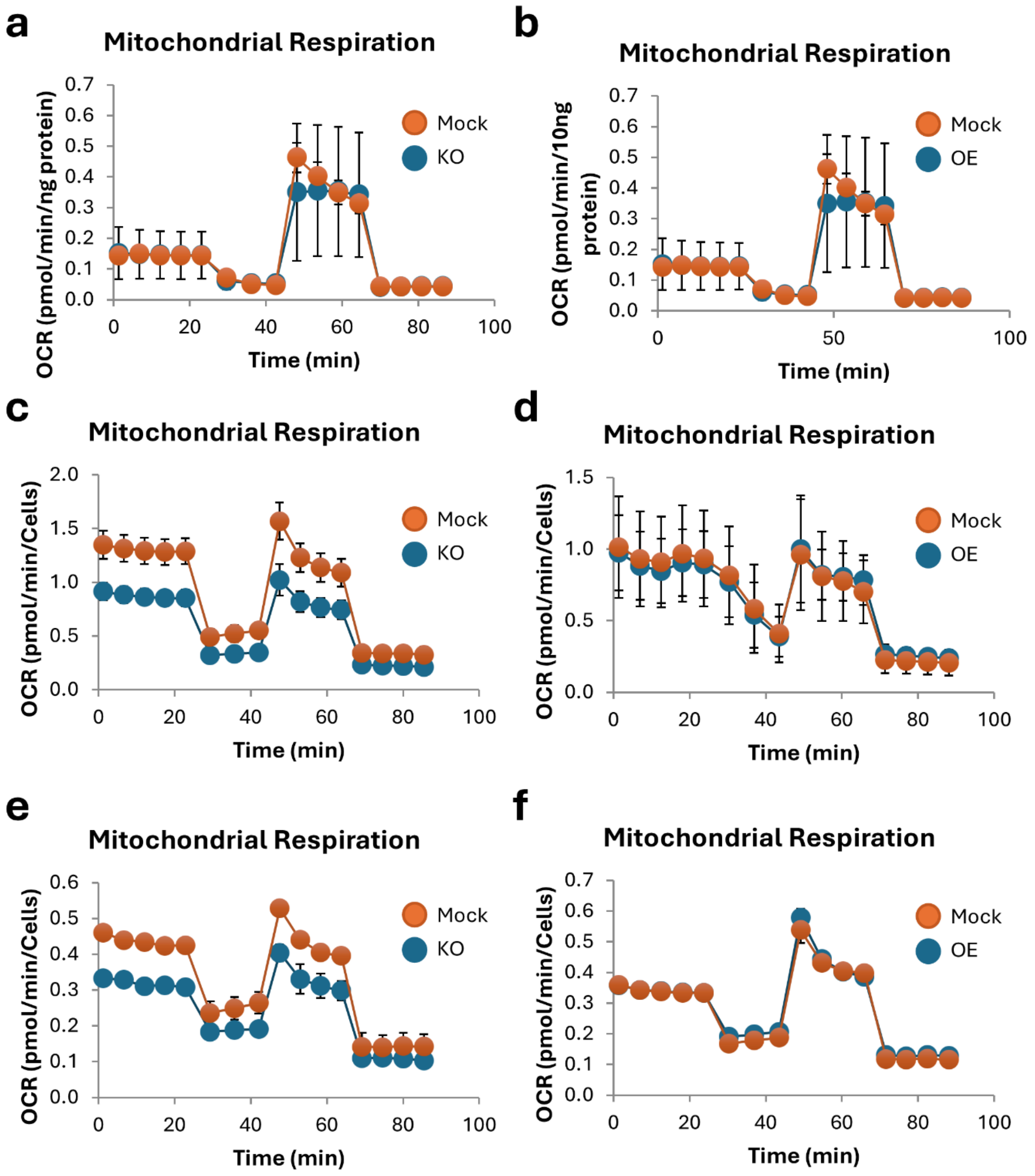
a
Mitochondrial Respiration
OCR (pmol/min/ng protein)
Time (min)
Mock
KO
b
Mitochondrial Respiration
OCR (pmol/min/10ng protein)
Time (min)
Mock
OE
c
Mitochondrial Respiration
OCR (pmol/min/Cells)
Time (min)
Mock
KO
d
Mitochondrial Respiration
OCR (pmol/min/Cells)
Time (min)
Mock
OE
e
Mitochondrial Respiration
OCR (pmol/min/Cells)
Time (min)
Mock
KO
f
Mitochondrial Respiration
OCR (pmol/min/Cells)
Time (min)
Mock
OE

**Supplementary figure 9:** Pre-ranked GSEA analysis of GOBP enrichment in the TE datasets.

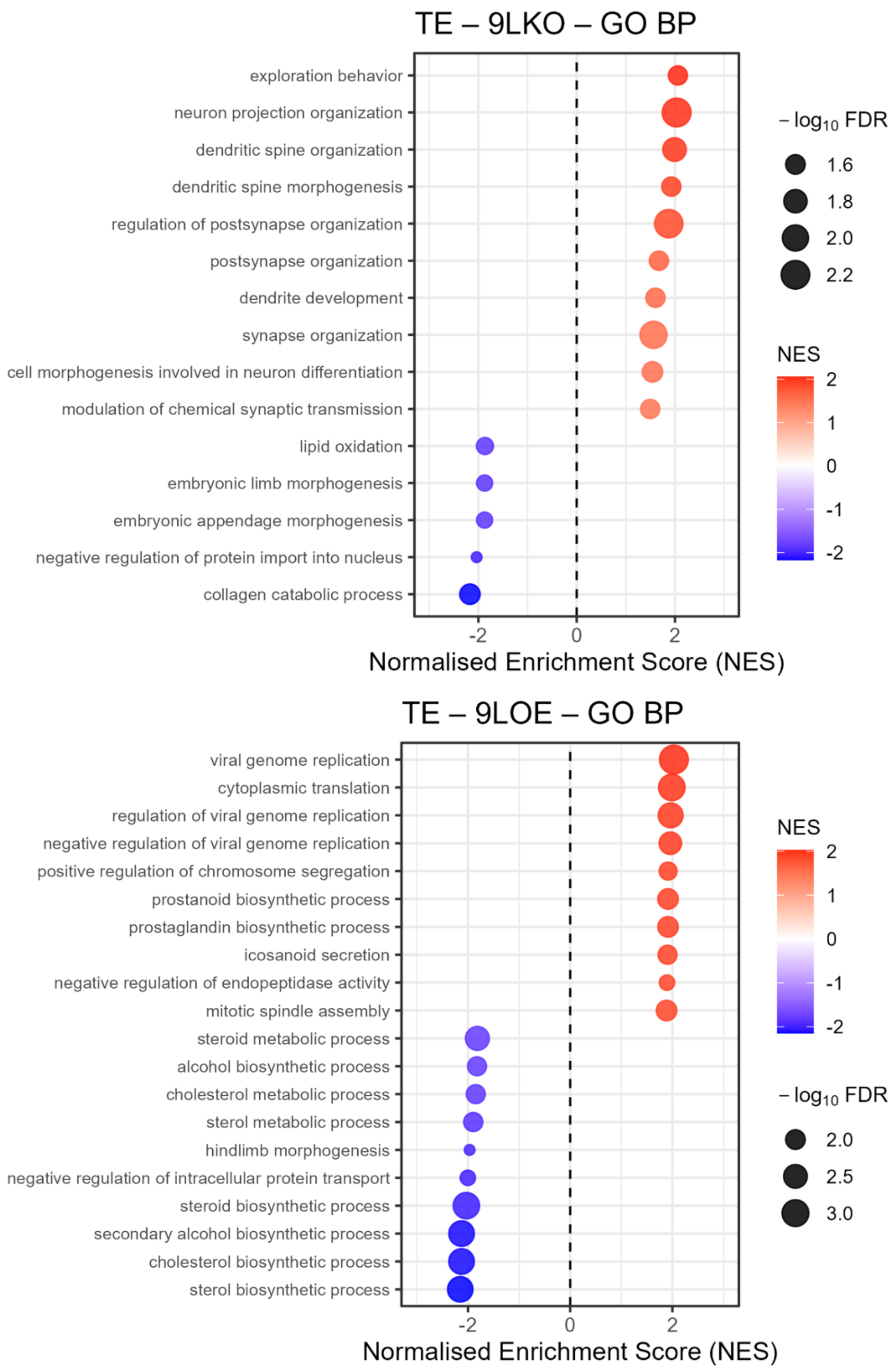

TE – 9LKO – GO BP
exploration behavior
neuron projection organization
dendritic spine organization
dendritic spine morphogenesis
regulation of postsynapse organization
postsynapse organization
dendrite development
synapse organization
cell morphogenesis involved in neuron differentiation
modulation of chemical synaptic transmission
lipid oxidation
embryonic limb morphogenesis
embryonic appendage morphogenesis
negative regulation of protein import into nucleus
collagen catabolic process
-2
0
2
Normalised Enrichment Score (NES)
$-\log_{10}$ FDR
1.6
1.8
2.0
2.2
NES
2
1
0
-1
-2
TE – 9LOE – GO BP
viral genome replication
cytoplasmic translation
regulation of viral genome replication
negative regulation of viral genome replication
positive regulation of chromosome segregation
prostanoid biosynthetic process
prostaglandin biosynthetic process
icosanoid secretion
negative regulation of endopeptidase activity
mitotic spindle assembly
steroid metabolic process
alcohol biosynthetic process
cholesterol metabolic process
sterol metabolic process
hindlimb morphogenesis
negative regulation of intracellular protein transport
steroid biosynthetic process
secondary alcohol biosynthetic process
cholesterol biosynthetic process
sterol biosynthetic process
-2
0
2
Normalised Enrichment Score (NES)
NES
2
1
0
-1
-2
$-\log_{10}$ FDR
2.0
2.5
3.0

**Supplementary figure 10:** ALKBH1 regulates global protein synthesis rates. **a:** Codon occupancy metagene plots across the coding sequences (CDS), downstream from the start codon, and upstream from the stop codon in KO (vs MockKO) and OE (vs MockOE) cells. **b:** Puromycin incorporation assay in transgenic GS-9L cells. Graph shows quantification of puromycin signal normalized to Actin-β. Asterisk: $p = 0.0002244$ on one-way ANOVA with Turkey's post hoc.

**a**

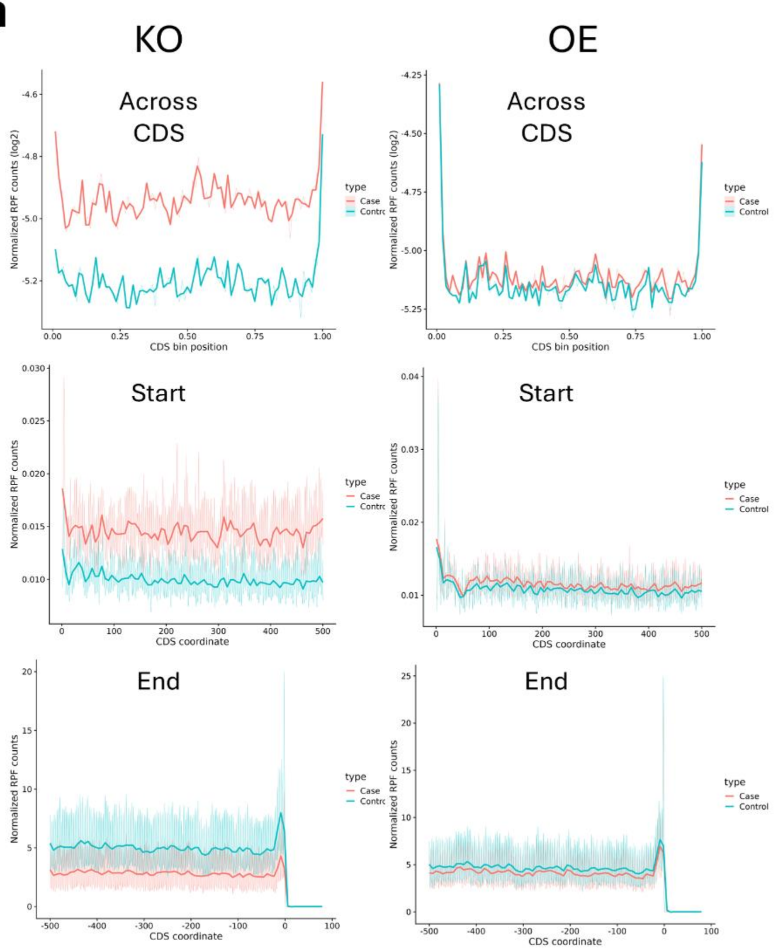
KO
OE
Across CDS
Start
End
Normalized RPF counts (log2)
Normalized RPF counts
CDS bin position
CDS coordinate
type
Case
Control

**b**

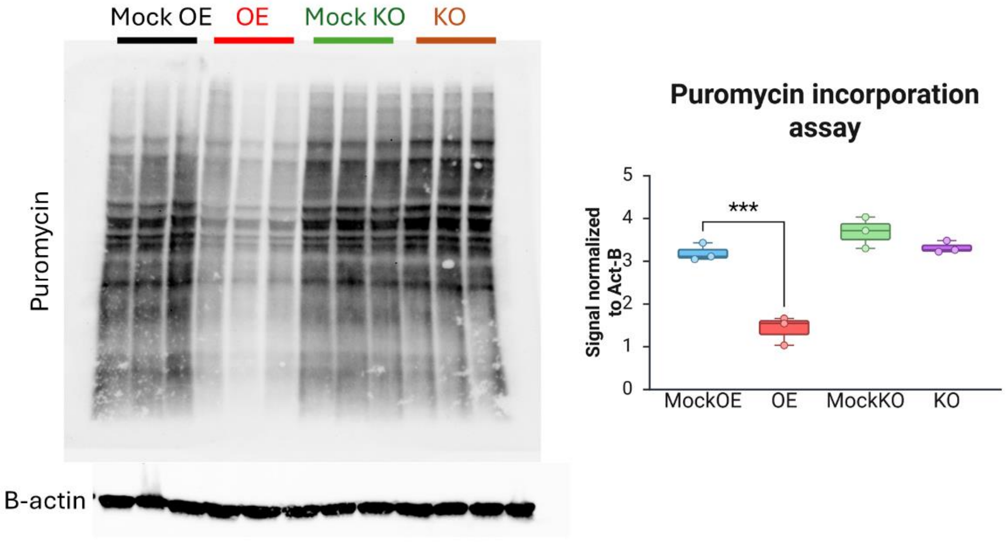
Mock OE
OE
Mock KO
KO
Puromycin
B-actin
Puromycin incorporation assay
Signal normalized to Act-B
***
MockOE
OE
MockKO
KO

**Supplementary figure 11: ALKBH1 modulates codon decoding. a:** Up and downregulated genes GC3 scores density plots in OE cells across datasets. **b:** Heatmap showing total codon usage of the up and downregulated genes in the Ribo-seq and TE datasets of the OE group.

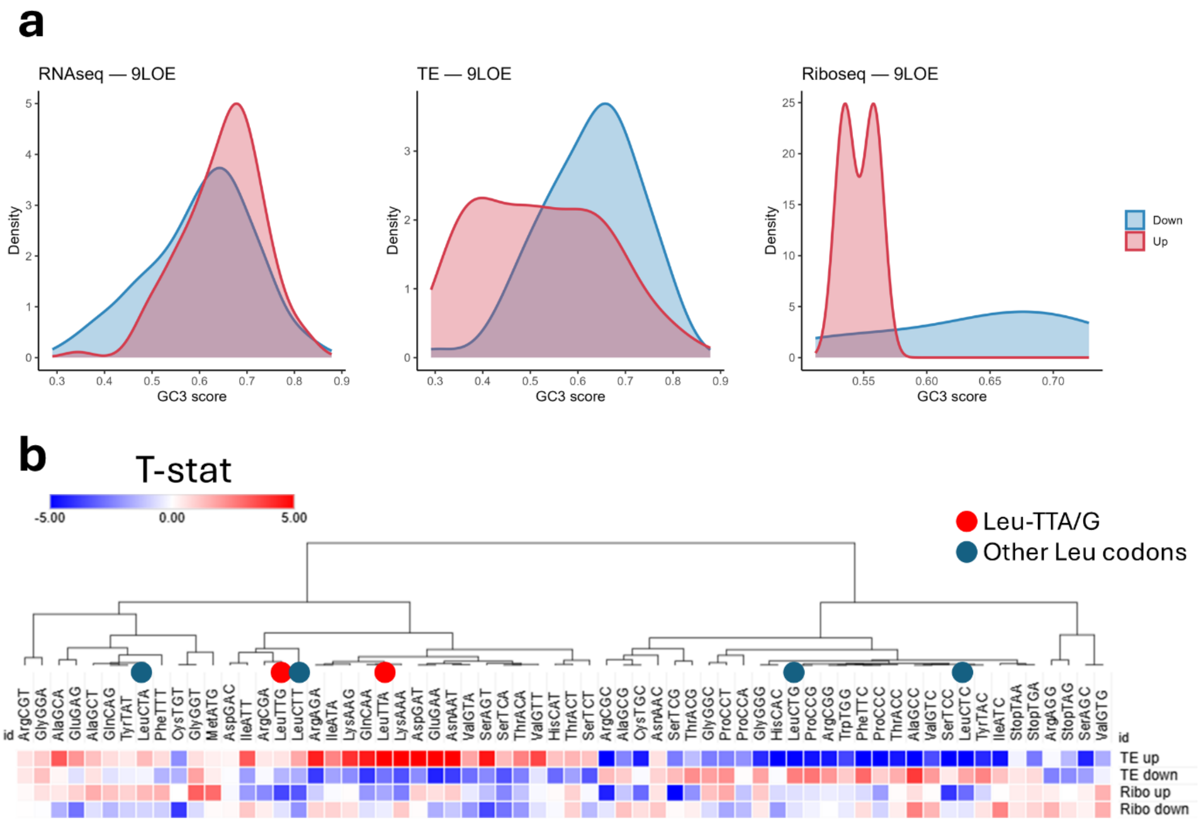

a
RNAseq — 9LOE
TE — 9LOE
Riboseq — 9LOE
Density
GC3 score
Down
Up
b
T-stat
-5.00
0.00
5.00
Leu-TTA/G
Other Leu codons
id
ArgCGT GlyGGA AlaGCA GluGAG AlaGCT GlnCAG TyrTAT LeuCTA PheTTT CysTGT GlyGGT MetATG AspGAC IleATT ArgCGA LeuTTG LeuCTT ArgAGA IleATA LysAAG GlnCAA LeuTTA LysAAA AspGAT GluGAA AsnAAT ValGTA SerAGT SerTCA ThrACA ValGTT HisCAT ThrACT SerTCT ArgCGC AlaGCG CysTGC AsnAAC SerTCG ThrACG GlyGGC ProCCT ProCCA GlyGGG HisCAC LeuCTG ProCCG ArgCGG TrpTGG PheTTC ProCCC ThrACC AlaGCC ValGTC SerTCC LeuCTC TyrTAC IleATC StopTAA StopTGA ArgAGG StopTAG SerAGC ValGTG
TE up
TE down
Ribo up
Ribo down

**Supplementary figure 12:** Leucine codon isoacceptors analysis: **a:** isoacceptors frequencies density plots of Leucine codons across the transcriptome showing which codon is more used (i.e., more optimal) in rat mRNAs. The same pattern is observed in human and mouse transcriptomes. **b:** Global heatmap of Leu isoacceptors ORA analysis. **c-e:** Dot plots showing the GOBP enrichment of the top 5% genes enriched in each designated Leu codon.

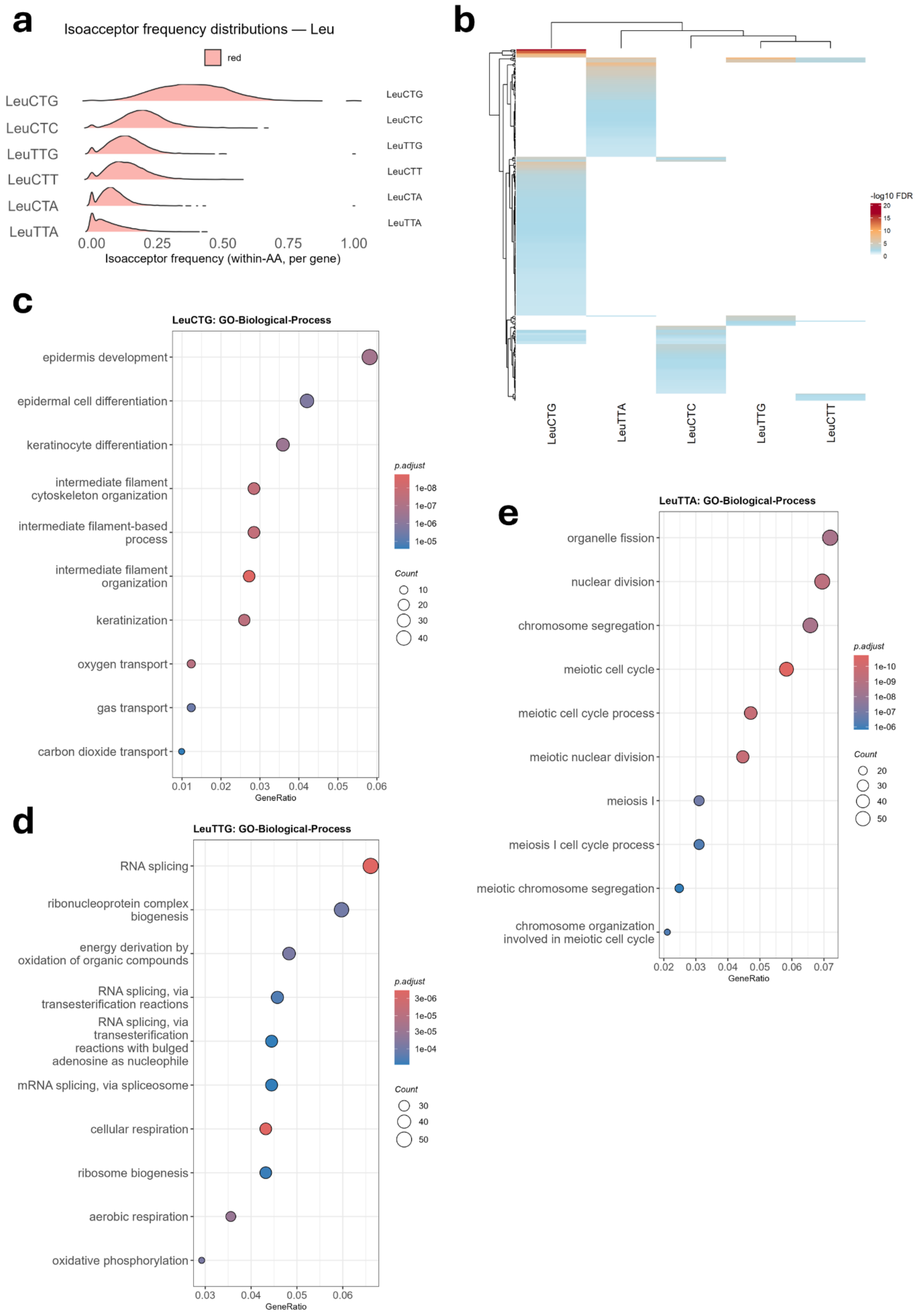
a
Isoacceptor frequency distributions — Leu
red
LeuCTG
LeuCTC
LeuTTG
LeuCTT
LeuCTA
LeuTTA
0.00
0.25
0.50
0.75
1.00
Isoacceptor frequency (within-AA, per gene)
b
-log10 FDR
20
15
10
5
0
LeuCTG
LeuTTA
LeuCTC
LeuTTG
LeuCTT
c
LeuCTG: GO-Biological-Process
epidermis development
epidermal cell differentiation
keratinocyte differentiation
intermediate filament cytoskeleton organization
intermediate filament-based process
intermediate filament organization
keratinization
oxygen transport
gas transport
carbon dioxide transport
0.01
0.02
0.03
0.04
0.05
0.06
GeneRatio
p.adjust
1e-08
1e-07
1e-06
1e-05
Count
10
20
30
40
d
LeuTTG: GO-Biological-Process
RNA splicing
ribonucleoprotein complex biogenesis
energy derivation by oxidation of organic compounds
RNA splicing, via transesterification reactions
RNA splicing, via transesterification reactions with bulged adenosine as nucleophile
mRNA splicing, via spliceosome
cellular respiration
ribosome biogenesis
aerobic respiration
oxidative phosphorylation
0.03
0.04
0.05
0.06
GeneRatio
p.adjust
3e-06
1e-05
3e-05
1e-04
Count
30
40
50
e
LeuTTA: GO-Biological-Process
organelle fission
nuclear division
chromosome segregation
meiotic cell cycle
meiotic cell cycle process
meiotic nuclear division
meiosis I
meiosis I cell cycle process
meiotic chromosome segregation
chromosome organization involved in meiotic cell cycle
0.02
0.03
0.04
0.05
0.06
0.07
GeneRatio
p.adjust
1e-10
1e-09
1e-08
1e-07
1e-06
Count
20
30
40
50

**Supplementary figure 13:** Leucine codon total codon analysis: **a:** Total codon frequencies density plots of Leucine codons across the transcriptome showing which codon is more used (i.e., more optimal) in rat mRNAs. **b:** Global heatmap of Leu total codons ORA analysis. **c-e:** Dot plots showing the GOBP enrichment of the top 5% genes enriched in each designated Leu codon.

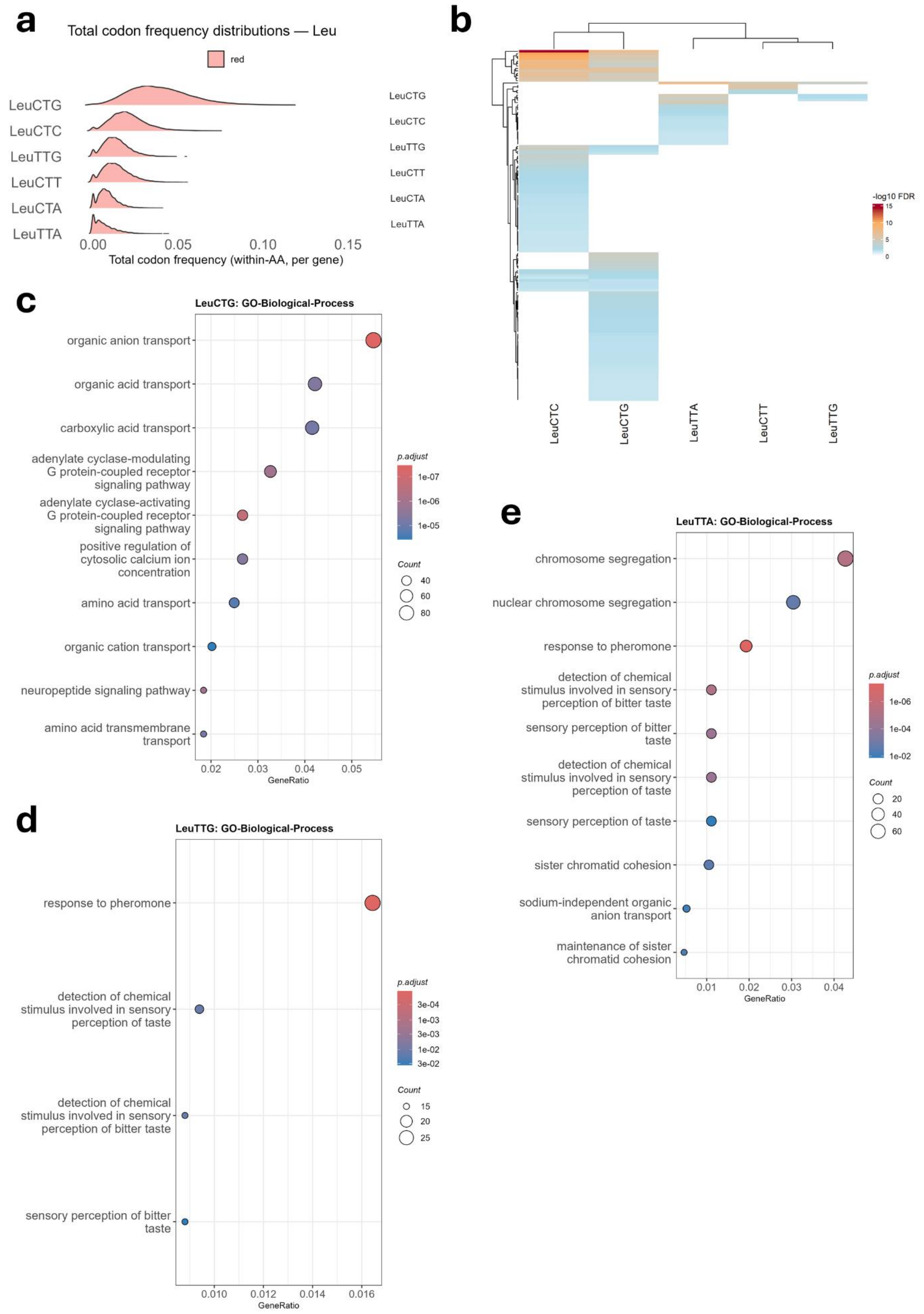
a
Total codon frequency distributions — Leu
red
LeuCTG
LeuCTC
LeuTTG
LeuCTT
LeuCTA
LeuTTA
0.00
0.05
0.10
0.15
Total codon frequency (within-AA, per gene)
b
-log10 FDR
LeuCTC
LeuCTG
LeuTTA
LeuCTT
LeuTTG
c
LeuCTG: GO-Biological-Process
organic anion transport
organic acid transport
carboxylic acid transport
adenylate cyclase-modulating G protein-coupled receptor signaling pathway
adenylate cyclase-activating G protein-coupled receptor signaling pathway
positive regulation of cytosolic calcium ion concentration
amino acid transport
organic cation transport
neuropeptide signaling pathway
amino acid transmembrane transport
GeneRatio
p.adjust
Count
d
LeuTTG: GO-Biological-Process
response to pheromone
detection of chemical stimulus involved in sensory perception of taste
detection of chemical stimulus involved in sensory perception of bitter taste
sensory perception of bitter taste
GeneRatio
e
LeuTTA: GO-Biological-Process
chromosome segregation
nuclear chromosome segregation
response to pheromone
detection of chemical stimulus involved in sensory perception of bitter taste
sensory perception of bitter taste
detection of chemical stimulus involved in sensory perception of taste
sensory perception of taste
sister chromatid cohesion
sodium-independent organic anion transport
maintenance of sister chromatid cohesion
GeneRatio

**Supplementary figure 14: a:** Heatmap showing the top 5 markers per cluster. **b:** Global cell type distribution across all conditions. **c:** Violin plot showing the expression of Gja1 across clusters. **d:** Violin plot showing the expression of Tenm2 across clusters.

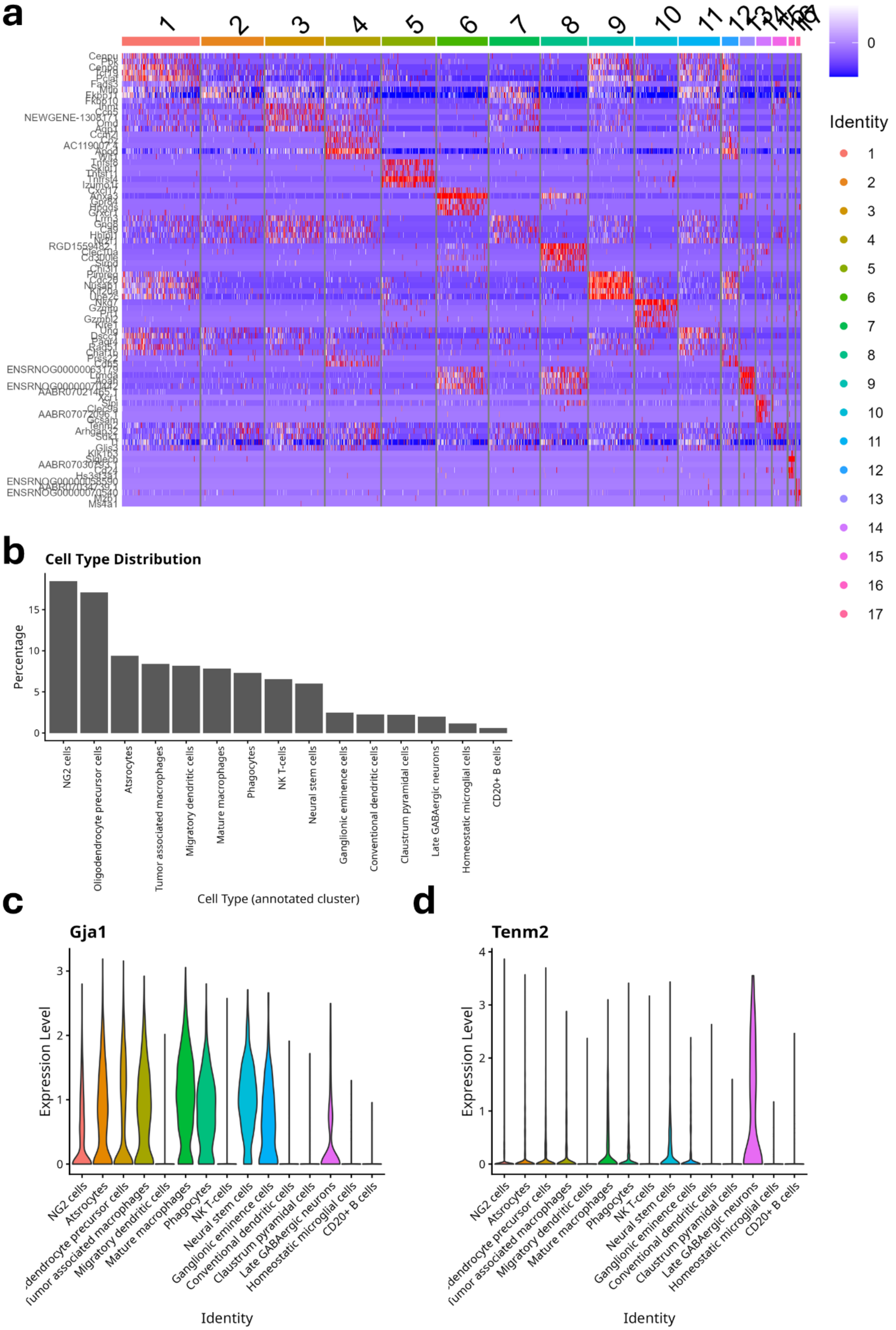
a
Identity
0
b
Cell Type Distribution
Percentage
0
5
10
15
NG2 cells
Oligodendrocyte precursor cells
Atsrocytes
Tumor associated macrophages
Migratory dendritic cells
Mature macrophages
Phagocytes
NK T-cells
Neural stem cells
Ganglionic eminence cells
Conventional dendritic cells
Claustrum pyramidal cells
Late GABAergic neurons
Homeostatic microglial cells
CD20+ B cells
Cell Type (annotated cluster)
c
Gja1
Expression Level
0
1
2
3
Identity
d
Tenm2
Expression Level
0
1
2
3
4
Identity

**Supplementary figure 15: a:** cell count per cluster by condition. **b:** Percentage of cells within each cluster by condition.

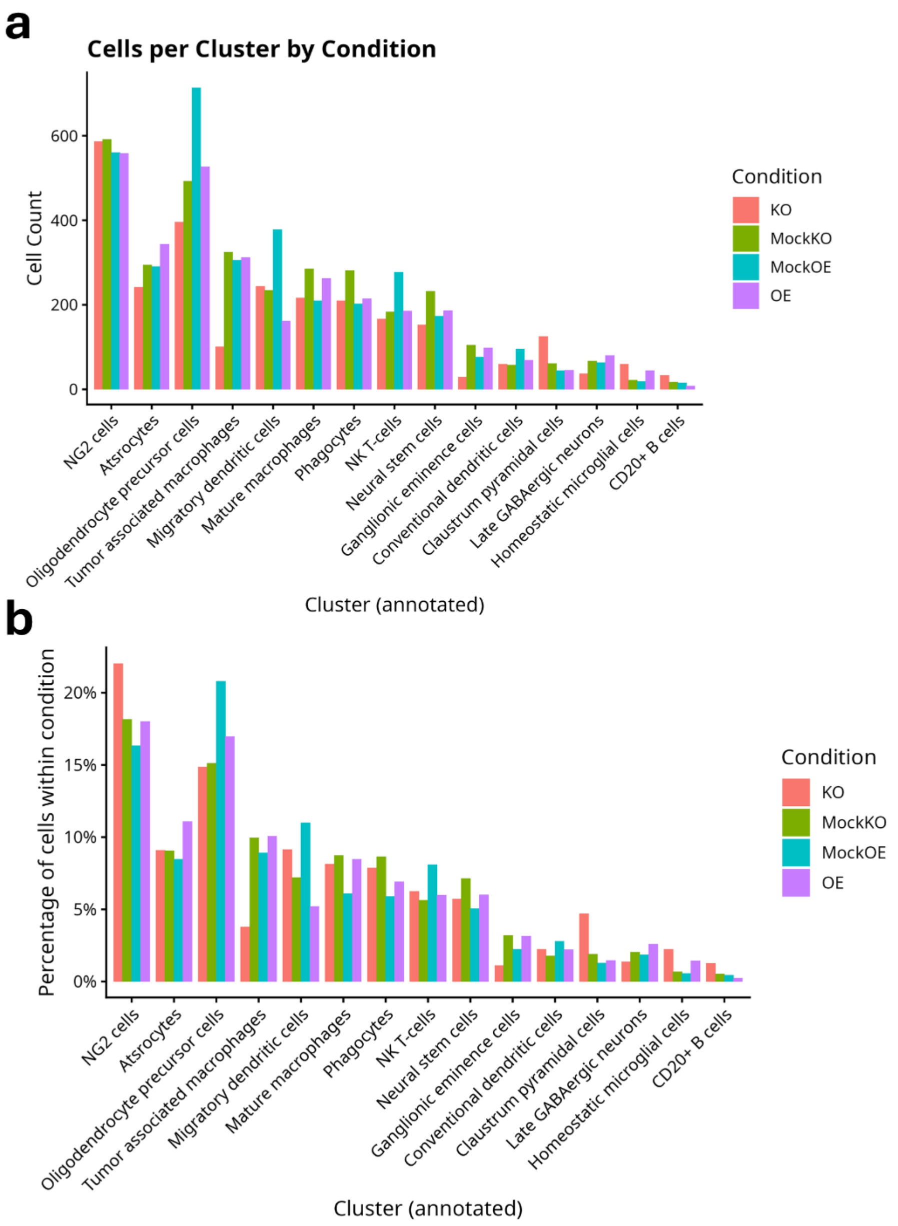
a
Cells per Cluster by Condition
Cell Count
0
200
400
600
Condition
KO
MockKO
MockOE
OE
NG2 cells
Atsrocytes
Oligodendrocyte precursor cells
Tumor associated macrophages
Migratory dendritic cells
Mature macrophages
Phagocytes
NK T-cells
Neural stem cells
Ganglionic eminence cells
Conventional dendritic cells
Claustrum pyramidal cells
Late GABAergic neurons
Homeostatic microglial cells
CD20+ B cells
Cluster (annotated)
b
Percentage of cells within condition
0%
5%
10%
15%
20%
Condition
KO
MockKO
MockOE
OE
NG2 cells
Atsrocytes
Oligodendrocyte precursor cells
Tumor associated macrophages
Migratory dendritic cells
Mature macrophages
Phagocytes
NK T-cells
Neural stem cells
Ganglionic eminence cells
Conventional dendritic cells
Claustrum pyramidal cells
Late GABAergic neurons
Homeostatic microglial cells
CD20+ B cells
Cluster (annotated)

**Supplementary figure 16:** Immunofluorescence staining of various proteins in tumors derived from different transgenic GS-9L cells. Bar size in panel **a** is 50µm and in panel **b** is 200µm.

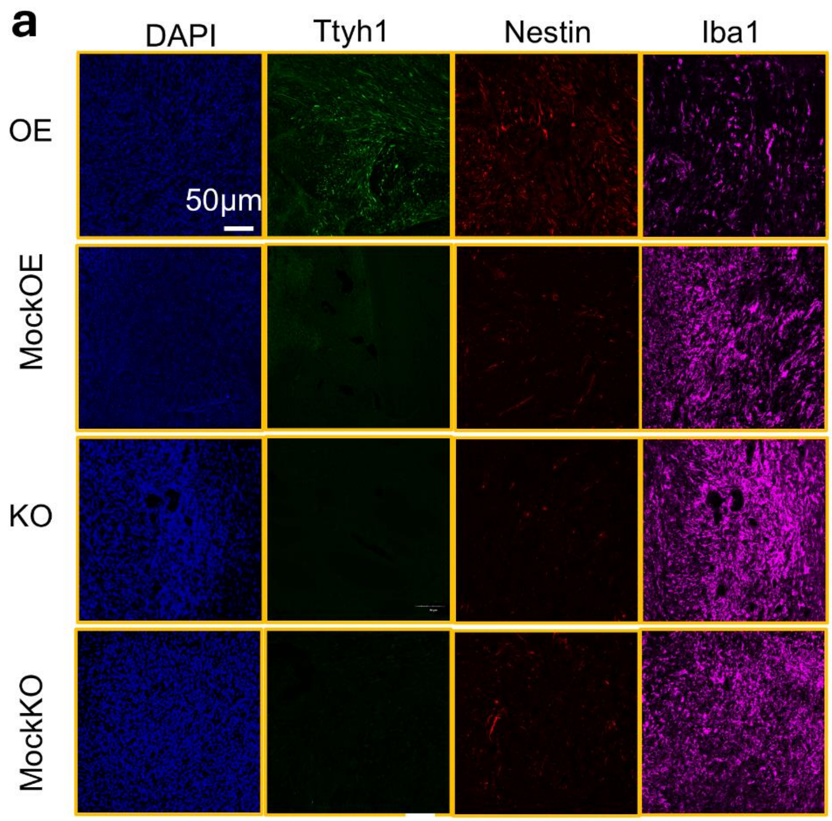
a
DAPI
Ttyh1
Nestin
Iba1
OE
MockOE
KO
MockKO
50μm

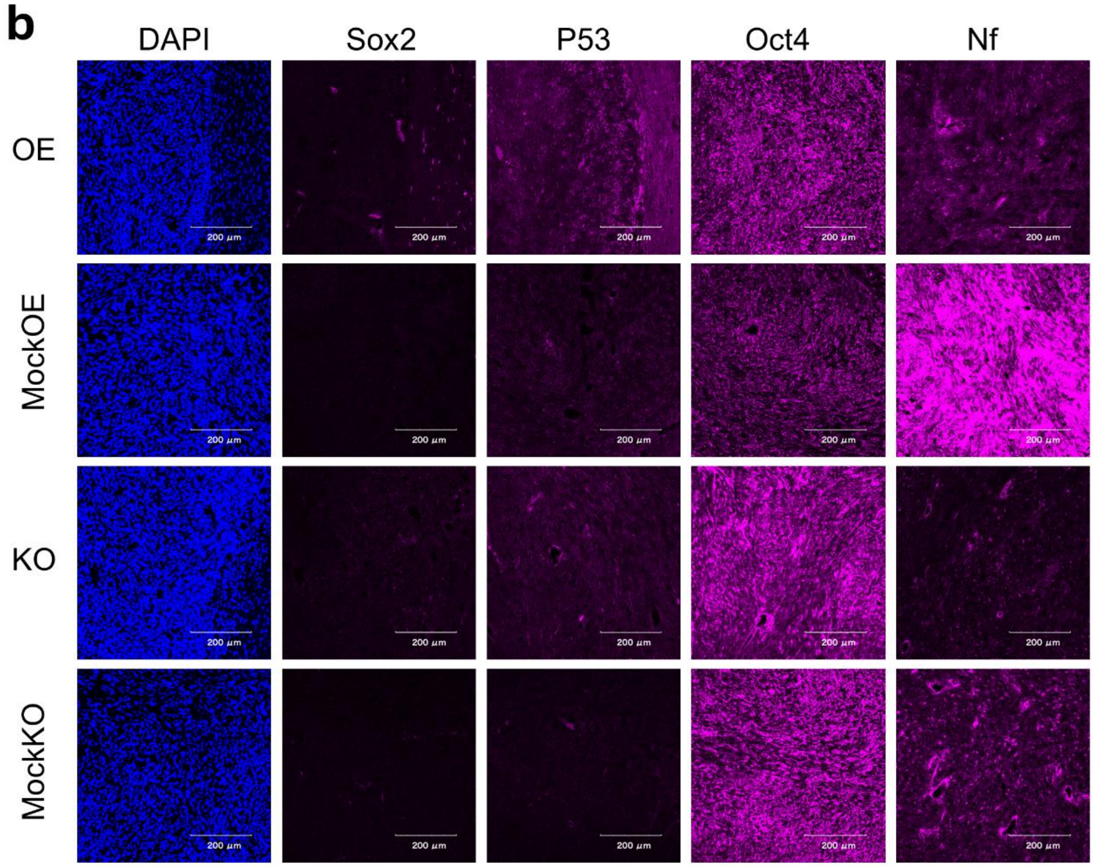
b
DAPI
Sox2
P53
Oct4
Nf
OE
MockOE
KO
MockKO

**Supplementary figure 17: a:** Number of interactions between cell clusters in KO vs Mock KO (Red lines indicate more interactions in KO and blue lines indicate more interactions in MockKO). **b:** Number of interactions between cell clusters in OE vs MockOE (Red lines indicate more interactions in OE and blue lines indicate more interactions in MockOE). **c:** Heatmap showing the pathway communication strength across conditions.

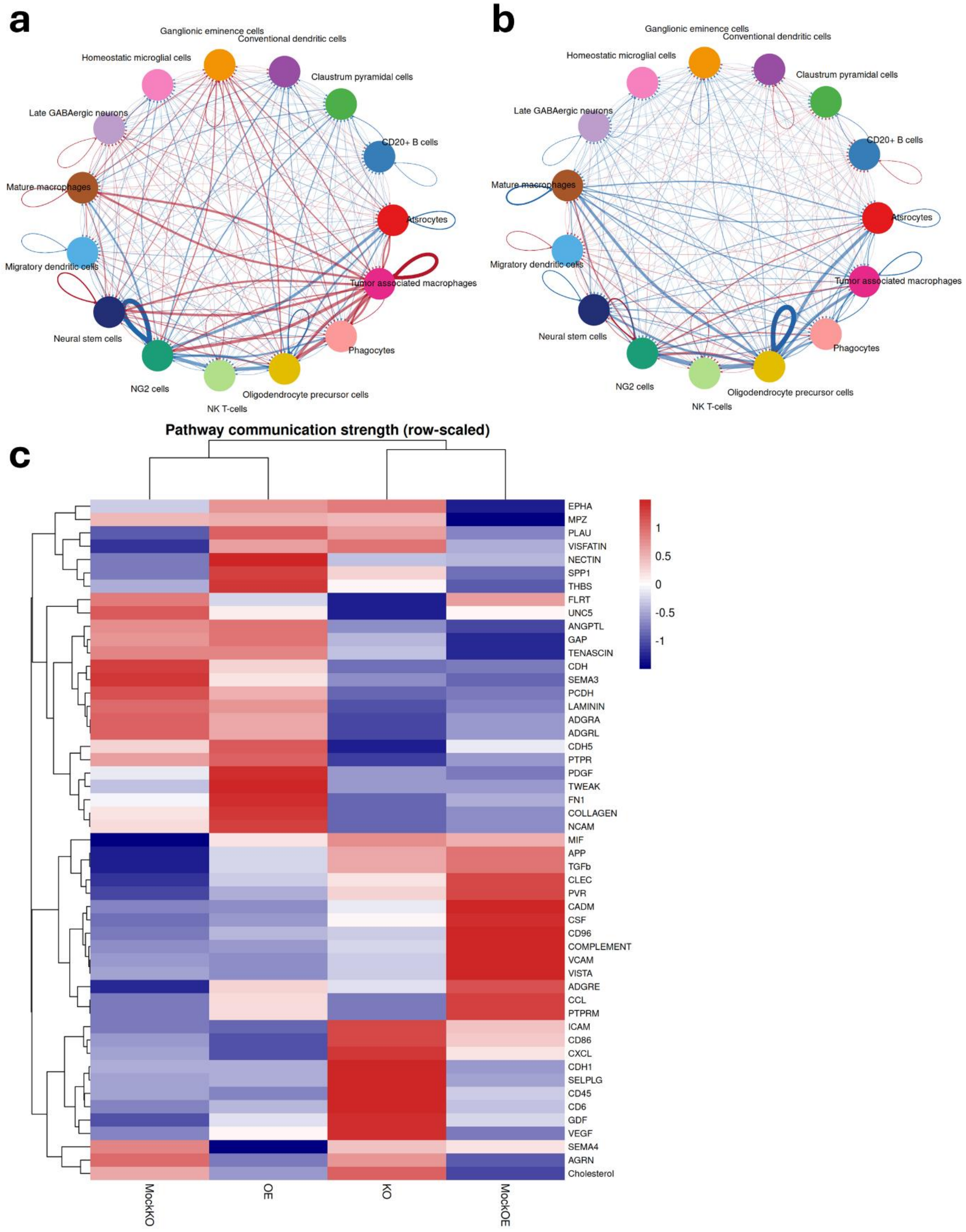
a
Ganglionic eminence cells
Conventional dendritic cells
Homeostatic microglial cells
Claustrum pyramidal cells
Late GABAergic neurons
CD20+ B cells
Mature macrophages
Atsrocytes
Migratory dendritic cells
Tumor associated macrophages
Neural stem cells
Phagocytes
NG2 cells
Oligodendrocyte precursor cells
NK T-cells
b
Ganglionic eminence cells
Conventional dendritic cells
Homeostatic microglial cells
Claustrum pyramidal cells
Late GABAergic neurons
CD20+ B cells
Mature macrophages
Atsrocytes
Migratory dendritic cells
Tumor associated macrophages
Neural stem cells
Phagocytes
NG2 cells
Oligodendrocyte precursor cells
NK T-cells
c
Pathway communication strength (row-scaled)
EPHA
MPZ
PLAU
VISFATIN
NECTIN
SPP1
THBS
FLRT
UNC5
ANGPTL
GAP
TENASCIN
CDH
SEMA3
PCDH
LAMININ
ADGRA
ADGRL
CDH5
PTPR
PDGF
TWEAK
FN1
COLLAGEN
NCAM
MIF
APP
TGFb
CLEC
PVR
CADM
CSF
CD96
COMPLEMENT
VCAM
VISTA
ADGRE
CCL
PTPRM
ICAM
CD86
CXCL
CDH1
SELPLG
CD45
CD6
GDF
VEGF
SEMA4
AGRN
Cholesterol
1
0.5
0
-0.5
-1
MockKO
OE
KO
MockOE

**Supplementary figure 18: a:** CellChat based analysis of Laminin pathway interactions and strength in each condition. **b:** CellChat based analysis of SPP1 pathway interactions and strength in each condition.

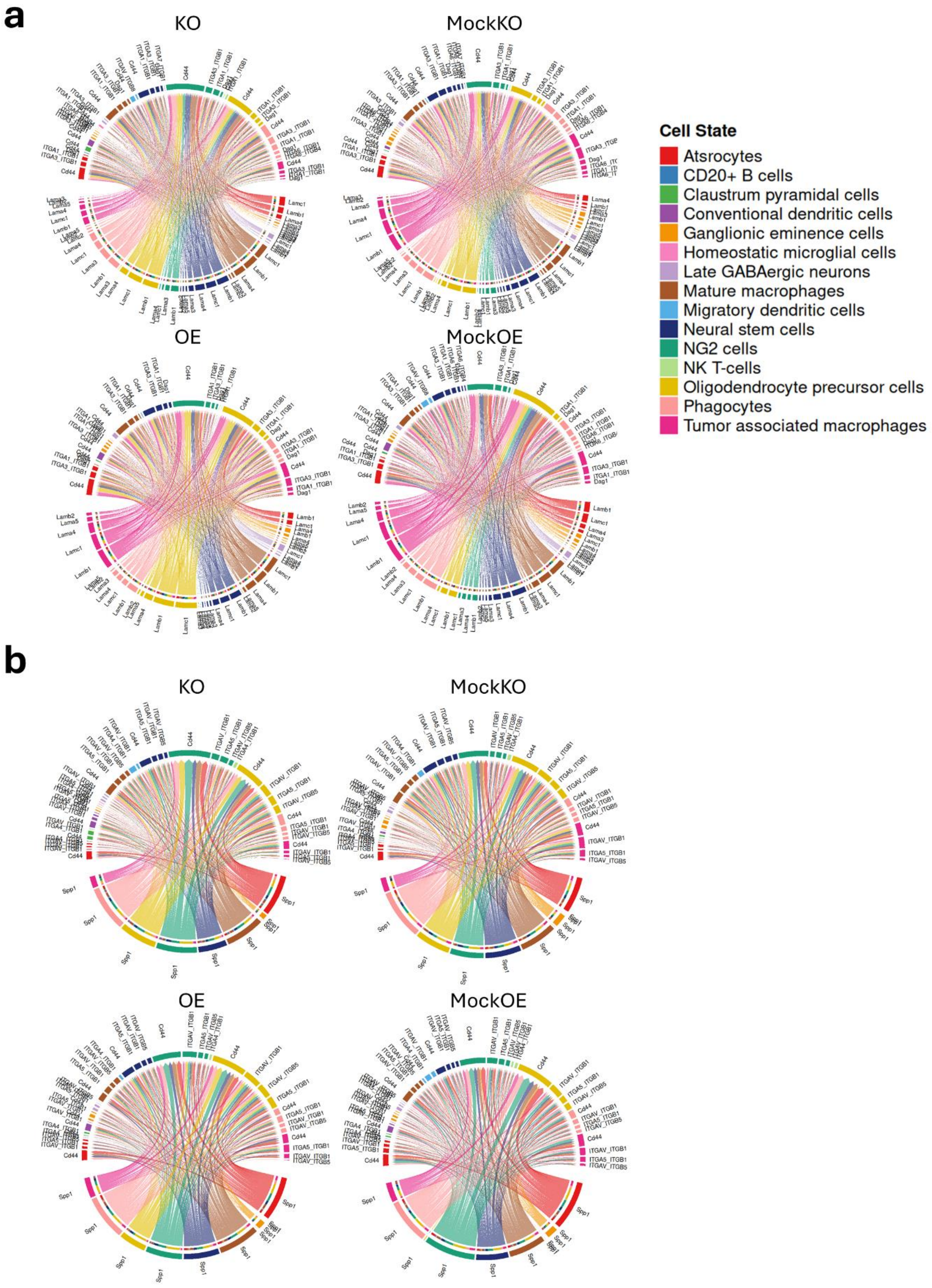
a
KO
MockKO
OE
MockOE
Cell State
Atsrocytes
CD20+ B cells
Claustrum pyramidal cells
Conventional dendritic cells
Ganglionic eminence cells
Homeostatic microglial cells
Late GABAergic neurons
Mature macrophages
Migratory dendritic cells
Neural stem cells
NG2 cells
NK T-cells
Oligodendrocyte precursor cells
Phagocytes
Tumor associated macrophages
b
KO
MockKO
OE
MockOE

**Supplementary figure 19: Differential gene expression analysis between conditions across clusters. a:** Volcano plot showing the differentially expressed genes between KO and MockKO in the OPC cluster. **b:** GOBP ORA analysis of significant genes in the KO vs MockKO comparison in the OPC cluster analysis. **c:** Volcano plot showing the differentially expressed genes between OE and MockOE in the OPC cluster. **d:** GOBP ORA analysis of significant genes in the OE vs MockOE comparison in the OPC cluster analysis. **e:** Volcano plot showing the differentially expressed genes between KO and MockKO in the NK T-cells cluster. **f:** Volcano plot showing the differentially expressed genes between OE and MockOE in the NK T-cells cluster. **g:** Volcano plot showing the differentially expressed genes between OE and MockOE in the NK T-cells cluster.

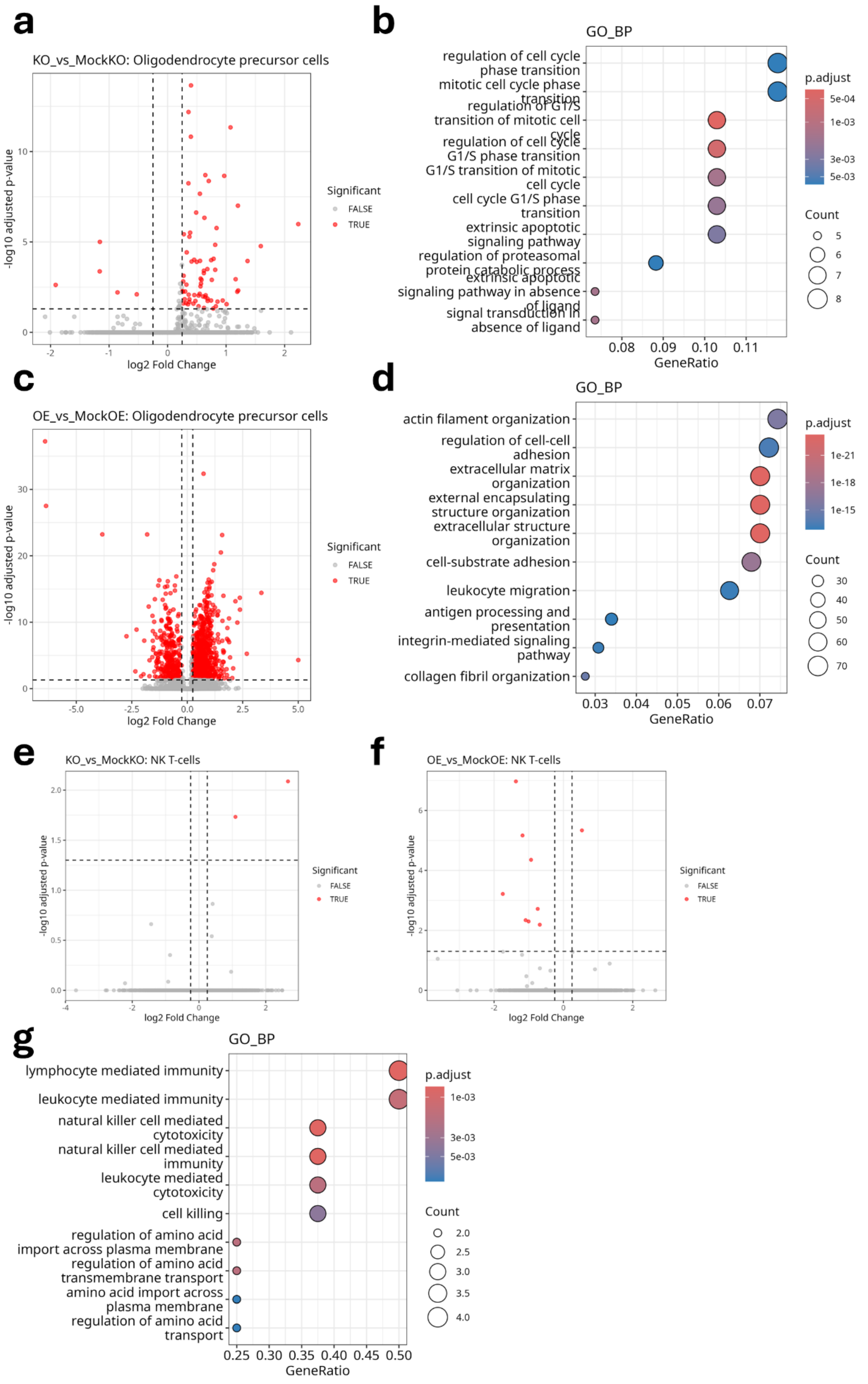
a
KO_vs_MockKO: Oligodendrocyte precursor cells
-log10 adjusted p-value
log2 Fold Change
Significant
FALSE
TRUE
b
GO_BP
regulation of cell cycle phase transition
mitotic cell cycle phase transition
regulation of G1/S transition of mitotic cell cycle
regulation of cell cycle G1/S phase transition
G1/S transition of mitotic cell cycle
cell cycle G1/S phase transition
extrinsic apoptotic signaling pathway
regulation of proteasomal protein catabolic process
extrinsic apoptotic signaling pathway in absence of ligand
signal transduction in absence of ligand
GeneRatio
p.adjust
Count
c
OE_vs_MockOE: Oligodendrocyte precursor cells
-log10 adjusted p-value
log2 Fold Change
Significant
FALSE
TRUE
d
GO_BP
actin filament organization
regulation of cell-cell adhesion
extracellular matrix organization
external encapsulating structure organization
extracellular structure organization
cell-substrate adhesion
leukocyte migration
antigen processing and presentation
integrin-mediated signaling pathway
collagen fibril organization
GeneRatio
p.adjust
Count
e
KO_vs_MockKO: NK T-cells
-log10 adjusted p-value
log2 Fold Change
Significant
FALSE
TRUE
f
OE_vs_MockOE: NK T-cells
-log10 adjusted p-value
log2 Fold Change
Significant
FALSE
TRUE
g
GO_BP
lymphocyte mediated immunity
leukocyte mediated immunity
natural killer cell mediated cytotoxicity
natural killer cell mediated immunity
leukocyte mediated cytotoxicity
cell killing
regulation of amino acid import across plasma membrane
regulation of amino acid transmembrane transport
amino acid import across plasma membrane
regulation of amino acid transport
GeneRatio
p.adjust
Count
